# Attractiveness in the eyes: A possibility of positive loop between transient pupil constriction and facial attraction

**DOI:** 10.1101/2020.04.02.021436

**Authors:** Hsin-I Liao, Makio Kashino, Shinsuke Shimojo

## Abstract

Contrary to the long-held belief of a close linkage between pupil dilation and attractiveness, we found an early and transient pupil constriction response when participants viewed an attractive face (and the effect of luminance/contrast is controlled). While human participants were making an attractiveness judgment on faces, their pupil constricted more for the more attractive (as-to-be-rated) faces. Further experiments showed that the effect of pupil constriction to attractiveness judgment extended to intrinsically aesthetic visual objects such as natural scene images (as well as faces) but not to line-drawing geometric figures. When participants were asked to judge the roundness of faces, pupil constriction still correlated with their attractiveness but not the roundness rating score, indicating the automaticity of the pupil constriction to attractiveness. When pupillary responses were manipulated implicitly by relative background luminance changes (from the pre-stimulus screen), the facial attractiveness ratings were in accordance with the amount of pupil constriction, which could not be explained solely by simultaneous or sequential luminance contrast. The overall results suggest that pupil constriction not only reflects but, as a part of self-monitoring and attribution mechanisms, also possibly contributes facial attractiveness implicitly.

## Introduction

Pupillary response not only reflects the peripheral nervous system’s activity in response to ambient luminance changes (i.e., the pupillary light reflex), but also the central nervous system’s activity underlying cognitive functions such as attention (Aston-Jones and Cohen, 2005; Einhäuser et al., 2008; Eldar et al., 2013), memory (Goldinger and Papesh, 2012; Naber et al., 2013; Zokaei et al., 2019), decision making (Einhäuser et al., 2010; de Gee et al., 2014), emotion (Partala and Surakka, 2003; Bradley et al., 2008), and interpersonal impressions and attitudes (Hess and Polt, 1960; Hess, 1965; Janisse, 1973; Hess, 1975). In the Middle Ages, women ingested belladonna to dilate their pupils, which was supposed to make them appear seductive. Nowadays, people can use cosmetic contact lenses to make the pupil appear larger (by changing the color and/or appearance of the iris). These cosmetic techniques are based on the long-held belief of a close link between pupil dilation and positive attitudes such as (sexual) interests and/or emotional arousal and thus of a mutual path between the actor and observer. Evidence in the early 60s showed that, actors’ faces with enlarged pupils were perceived as more attractive to observers (Hess, 1965; Stass and Willis, 1967; Hess, 1975; Bull and Shead, 1979). On the observer side, evidence indicated that people’s pupil dilated when they were viewing emotionally toned stimuli, such as pictures of a baby for female participants and pictures of a partially nude man or woman for female and male participants, respectively (Hess and Polt, 1960; Hess, 1965; cf. Janisse, 1973). This may be due to arousal and/or sexual attraction (Hess et al., 1965; Tombs and Silverman, 2004; Caryl et al., 2009; Rieger and Savin-Williams, 2012), which activates the sympathetic nervous system to induce pupil dilation. Together with activation of the mirror neuron system that may be involved in a positive circulation between the observer and the observed face (i.e., the actor), an intuitive prediction has been that the pupils of people who are attracted to faces they see dilate as an automatic response. Then, in turn, they would appear attractive to observers. Such interpersonal, positive feedback has been assumed for a long time.

However, there is room for skepticism because the dynamic of the pupillary response to attractiveness could be more complicated than has been thought. For example, the pupillary dilation (in observers) found in the early era may have been confounded with stimulus luminance or contrast to which the pupil responds most sensitively and/or insufficient baseline conditions (Janisse, 1973). Recent studies, which have had finer control over stimulus luminance and contrast with various tested conditions, have found that the pupil dilates to not only positive but also to negative emotional stimuli (Bradley et al., 2008; Burley et al., 2018). This suggests that it dilates to arousal stimuli in general, not particularly to a positive emotion and/or evaluation such as attractiveness. Moreover, most evidence from previous studies was based on pupil size averaged over several seconds, while the participants were asked to just passively view the stimuli (e.g., about 10 s in Hess, 1965; Nunnally et al., 1967; Stass and Willis, 1967; Koff and Hawkes, 1968; Barlow, 1969; Atwood and Howell, 1971; Rieger and Savin-Williams, 2012; Attard-Johnson et al., 2019; 2–6 s in Bradley et al., 2008). The long-lasting, sustained pupil dilation response reflecting arousal may be different from the fast, transient component which presumably reflects other cognitive states and thus affect the feeling of attractiveness. Indeed, other studies showed that pupils in general quickly constrict in response to the mere onset of visual presentation (even when the mean luminance is equated, e.g., Kimura et al. (2014) and that this early and reflexive pupillary constriction response is modulated by various cognitive factors such as memory (Naber et al., 2013), attention (Binda et al., 2013; Mathôt et al., 2013; Binda et al., 2014; Mathôt et al., 2014), and perceptual brightness when the physical luminance is kept the same (Laeng and Endestad, 2012; Suzuki et al., 2019). For instance, in Naber et al. (2013), participants were asked to memorize various natural scene images presented one by one (memorization phase) to recall later in the retrieval phase. The results showed that during the memorization phase, pupils constricted more strongly to certain images, which, upon retrieval were found to be better memorized. Taking into the evidence that people tend to better memorize attractive faces than they do moderately attractive ones (Shepherd and Ellis, 1973), we hypothesize that the pupil constricts more strongly for more attractive faces, at least during the encoding and/or memorization period, although the underlying mechanism remains unclear.

Aside from the literature on pupil responses to attractiveness, the issues can be discussed in a different context, namely affective decision making, which is a dynamic process to which various factors contribute, such as physiological arousal (e.g., the somatic marker hypothesis, Damasio, 1996), gaze (Shimojo et al., 2003), and perceptual fluency via mere exposure (Zajonc, 1968). Shimojo and colleagues demonstrated that active gaze engagement not only reflects but also affects preference decision making (the “gaze cascade” effect), suggesting a positive loop between seeing and liking (Shimojo et al., 2003). They simply revealed a gaze bias towards a to-be-chosen face to show that gaze *reflects* preference, but they were also successful in biasing preference decisions by manipulating gaze, thus demonstrating that gaze also *affects* preference. Just as gaze allows foveal scrutiny, pupil constriction improves visual acuity (Campbell and Gregory, 1960). Thus, we hypothesize that pupil constriction may also be actively involved in, or at least concur with the formation of preference via an enhancement of seeing and thus liking.

Along that line, we further speculate that the more implicit the information (causal factor) is, the stronger the decision-making processing may be affected. This seemingly counter-intuitive prediction is proved true at least occasionally in the literature regarding to the mere exposure effect (Bornstein, 1989). Under certain conditions, repetitively presented stimuli get more preferable (i.e., a stronger mere exposure effect is produced) when they are presented subliminally rather than suprathreshold. This pattern of results has been interpreted to mean that when information is implicit, i.e., subliminal, participants often do not causally attribute their decision to the repetitive presented stimuli per se and are thus more likely to attribute it to their own internal preference. The misattribution in affective decision-making was observed in the “gaze cascade effect” mentioned above. When people’s preferences were affected by their gaze manipulated (Shimojo et al. 2003, Exp. 2), most of them were not aware of the gaze bias to begin with, and those few who were aware of it did not attribute their preference to it. There was yet another study in which the participants were fully aware of all the stimuli (again faces); however, they confused their intended choice with the actual outcome. That is, they thought they preferred a particular face but in fact chose a different one beforehand, which is known as choice blindness (Johansson et al., 2005). In such cases, the retrospectively derived reasons for why a choice is made are inevitably the result of misattribution. The influence of pupil constriction on attractiveness judgment, or the concurrence of them, if it occurs, could also be misattributed and implicit. This is because the pupillary response itself is implicit (more so than gaze shifts) and thus cannot be voluntarily controlled or attentively introspected.

## Materials and Methods

We conducted seven experiments to address the issue of pupillary response and attractiveness judgment. Experiments 1 and 2 examined how the pupil responded when seeing an image that was evaluated as attractive. Experiments 3, 4 and 5 were conducted, all together with Experiments 1 and 2, to examine potential factors that might affect the result and/or account for the inconsistency between our finding (pupil constricts to attractiveness) and the literature, which demonstrated pupil dilation to attractiveness. The factors included stimulus presentation time, task demand (attractiveness judgment, roundness judgment, or passive viewing), stimulus category (faces, natural scenes, or geometric figures), and overall pupil response pattern (constriction or dilation) caused by sequential luminance contrast change. Experiment 6 examined whether implicit pupil manipulation contributes to attractiveness judgment, and Experiment 7 ruled out potential confounding in Experiment 6. Table 1 shows the overview of the critical manipulations in each experiment.

**Table 1.** Overview of the critical manipulations in each experiment.

|  | Objectives | Procedure | Task | Sti. | Overall pupil response pattern caused by sequential luminance contrast change |
| --- | --- | --- | --- | --- | --- |
| E1 | To examine how the pupil responds to attractiveness. | Participants made the judgment of an image while their pupillary responses were recorded. | Att., Rnd. | FC, GF | Pupil constriction to FC, Pupil dilation to GF |
| E2 | To replicate the result of E1 by using equal luminance images. | Same as E1 | Att., Rnd. | FC (eq), NS (eq) | Pupil constriction to both FC and NS (even when the mean luminance change is equated) |
| E3 | To examine whether stimulus presentation duration affects the result. | Participants viewed the image for 5 s and held their response after the stimulus presentation. | Att. | FC (eq) | Pupil constriction to FC (even when the mean luminance change is equated) |
| E4 | (1) To examine whether task demand affects the result.<br>(2) To minimize the effect of stimulus luminance on pupils. | Same as E3, or just viewing the stimulus for 5 s without judgment | Att., Pas. | FC (line) | Pupil constriction to FC (slightly) |
| E5 | To examine whether overall pupil response pattern affects the result . | Same as E1 | Att. | FC, GF | Pupil dilation to FC, Pupil constriction to GF |
| E6 | To examine whether implicit pupil size manipulation contributes to attractiveness. | Same as E1 | Att. | FC | The amount of pupil size change was manipulated. |
| E7 | To rule out the confounding of sequential luminance contrast in E6 | Same as E1 | Att. | FC | The amount of pupil size change was unchanged, while the sequential luminance contrast change was remained. |

Att. = Attractiveness task, Rnd. = Roundness task, Pas. = Passive viewing, Sti. = Stimuli, FC = Faces, GF = Geometric Figures, NS = Natural Scenes, eql = equal luminance, line = line drawing

In Experiment 1, participants looked at a face presented at the center of a screen and rated how attractive the face was on a scale from 1 (least attractive) to 9 (most attractive) while an infrared camera recorded their pupillary responses. Data were sorted based on the attractiveness judgment individually to examine how the pupil reacted to attractive faces (the face-attractiveness condition). Two other conditions were added to examine whether the effect of pupil constriction to attractive faces, if it occurs, was specific to faces or attractiveness judgment. In the geometric figure-attractiveness condition, participants evaluated the attractiveness of a geometric figure, but not a face. In the face-roundness condition, they viewed the same set of the faces to rate how round the faces were, ignoring their attractiveness. The three conditions were conducted in separate blocks in a counterbalanced order across participants. Each trial started with a 3-s gray fixation display, followed by the target image (faces or geometric figures). Participants were free to inspect the stimuli as long as they wanted before making a decision.

Experiment 2 aimed to replicate the finding of pupil constriction to attractive faces with additional luminance controls. First, the luminance among the faces was equated, and the mean luminance of the faces, as well as that of the fixation display presented before the faces was the same as the background (so that there was no mean luminance change over time). Second, instead of using line-drawing geometric figures, we used natural scenes. The photos were image processed to equate their mean luminance by following the same procedure as for the face images. The rest of the experimental procedures were the same as in Experiment 1.

Experiment 3 aimed to examine the time course of the effect of pupil constriction to attractiveness. In other words, does the effect occur only shortly after the stimulus presentation or does it last long for several seconds? In experimental procedure, the face image was presented for 5 s and participants were asked to hold their judgment response until the face disappeared. Their pupillary responses were recorded during the whole 5-s inspection period.

Experiment 4 had two purposes. First, we aimed to re-examine the effect of pupil constriction to attractive faces over time as in Experiment 3 by adding another factor: task demand. This was to examine whether pupil dilation to attractiveness could be observed in later time course and when no task demand was involved, as in the similar condition where pupil dilation was observed in the literature. To this end, in addition to the attractiveness judgment condition as was done in Experiment 3, we added a passive viewing condition, in which participants viewed the same set of images presented in the attractiveness judgment condition without any task demand. The two conditions were conducted in counterbalanced order across participants. Pupil data in the passive viewing condition were sorted by attractiveness ratings obtained in the attractiveness judgment condition for each participant individually. The second purpose of Experiment 4 was to minimize the effect of stimulus luminance change on pupils. We therefore used line-drawing face images. The faces were presented with their luminance contrast slowly enhanced, instead of sudden flash as in the previous experiments, to minimize the transient change on the screen.

In Experiment 5, we examined whether the baseline pupil response (constriction or dilation) is critical to the effect of pupil constriction to attractiveness. The luminance of the fixation display prior to the target display, as well as the background of the target display, was manipulated so that the pupil response baseline change was dilation to faces and constriction to geometric figures, in the opposite direction compared with Experiment 1.

Experiment 6 was to examine whether pupil constriction contributes to facial attractiveness judgment. To this end, while keeping the target image the same, we manipulated the luminance of the fixation display (prior to the target display) to so that it would change from black or gray to alter the amount of pupil constriction when the page flipped. Due to the nature of the pupillary light reflex (Ellis, 1981), the pupil should constrict more strongly when the target image follows a black than a gray fixation display. We also changed the luminance of the target background to black or gray, to serve as fillers to make the critical manipulation, i.e., the fixation display change, less noticeable. With this manipulation, we also aimed to examine the relative contribution of pupil constriction and simultaneous luminance contrast (induced by the target background) to attractiveness judgment. Although the luminance of the target background may also affect the pupillary response, its influence is expected to be smaller than that of the fixation display. If the attractiveness judgment is affected more by the simultaneous contrast than the pupil constriction, the target background luminance should have a stronger influence on attractiveness judgment than the pre-stimulus fixation display. Participants rated the attractiveness of the faces presented at the center of the target display as in Experiment 1 (Figure 6A).

**Figure 1.**
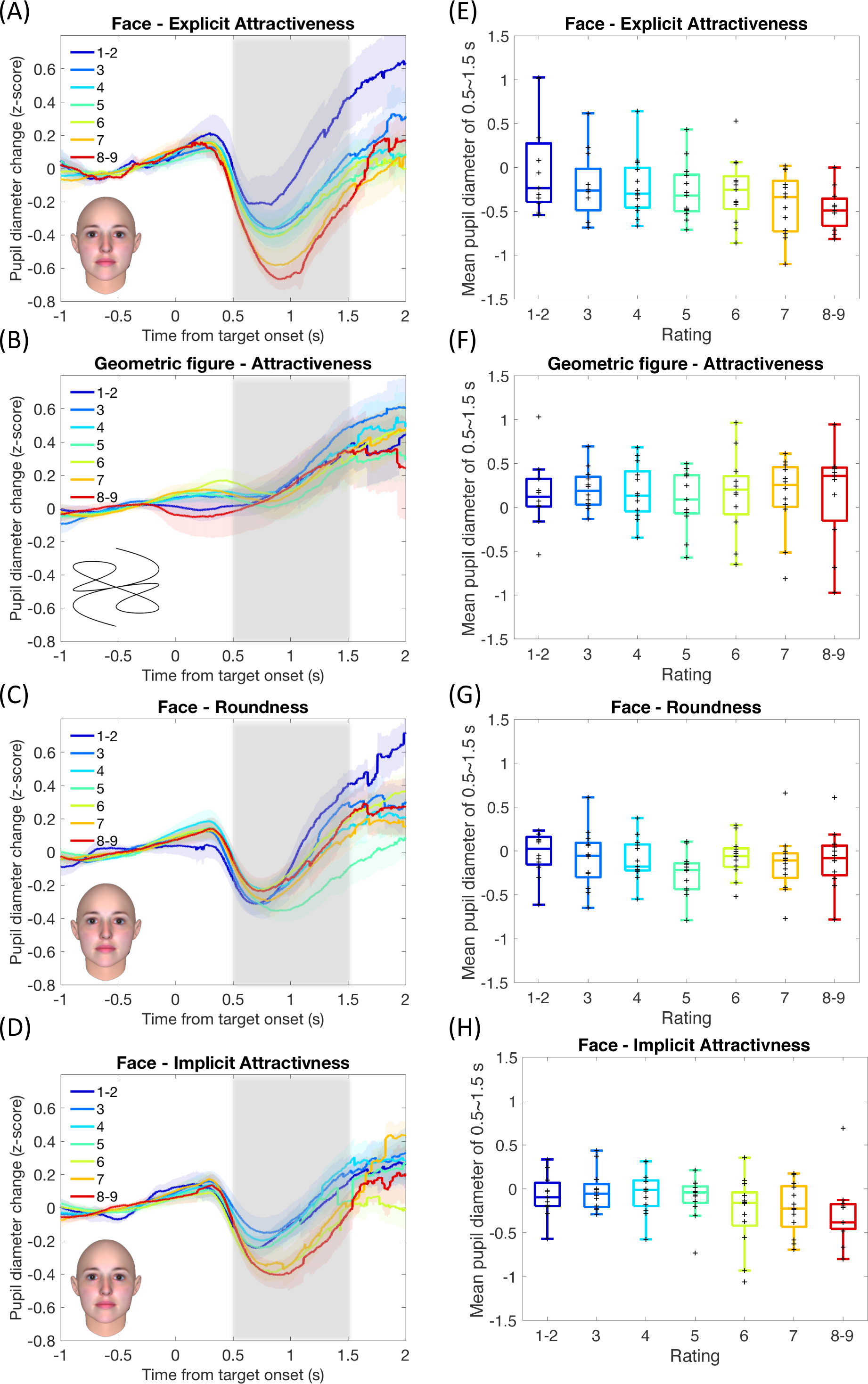
Pupil response results in Experiment 1. Sample stimulus images are shown corresponding to individual conditions. (A)–(D) Mean pupil diameter change as a function of time reference to the target onset during (A) attractiveness judgment for faces, (B) attractiveness judgment for geometric figures, (C) roundness judgment for faces, and (D) roundness judgment for faces when the data was sorted by the attractiveness of the faces. Curves are parameterized with rating score depending on individual participants’ choices (1 for least attractive and 9 for most attractive for panel A, B, and D; 1 for least round and 9 for roundest for panel C). The color shadows represent standard errors among participants. The gray shadow represents the time window for averaging the pupil size to present the amount of pupil constriction for statistical analysis (see Materials and Methods for details and Table 3 for results). (E)–(H) Box plots of the mean pupil size over the specified time window. The plus signs represent individual data.

**Figure 2.**
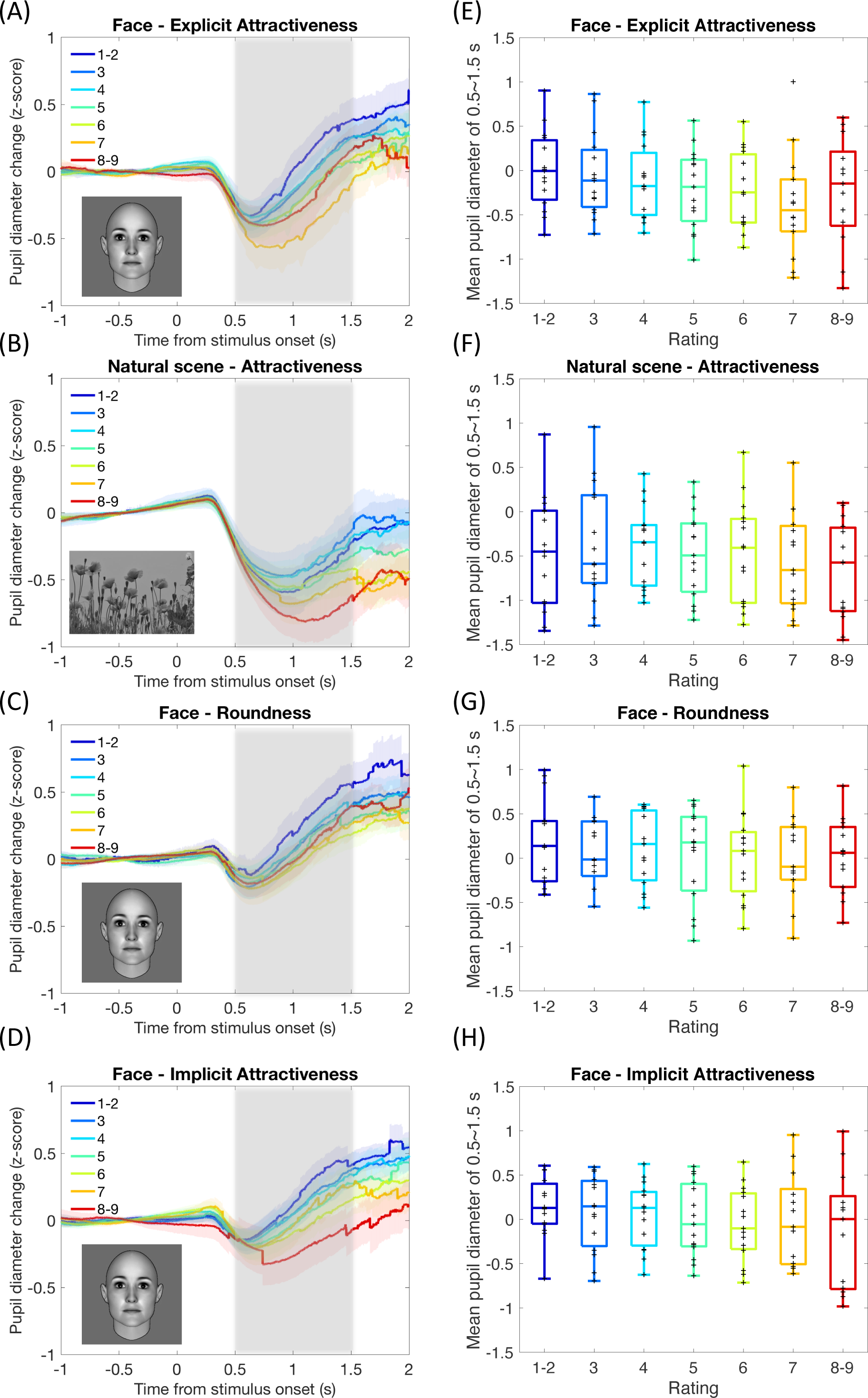
Pupil response results in Experiment 2. Sample stimulus images (luminance equated) are shown corresponding to individual conditions. (A)–(D) Mean pupil diameter change as a function of time reference to the target onset during (A) attractiveness judgment for faces, (B) attractiveness judgment for natural scenes, (C) roundness judgment for faces, and (D) roundness judgment faces when the data was sorted by the attractiveness of the faces. Curves are parameterized with rating score depending on individual participants’ choices (1 for least attractive and 9 for most attractive for panel A, B, and D; 1 for least round and 9 for roundest for panel C). The color shadows represent standard errors among participants. The gray shadow represents the time window for averaging the pupil size to present the amount of pupil constriction, for statistical analysis (see Materials and Methods for details and Table 3 for results). (E)–(H) Box plots of the mean pupil size over the specified time window. The plus signs represent individual data.

**Figure 3.**
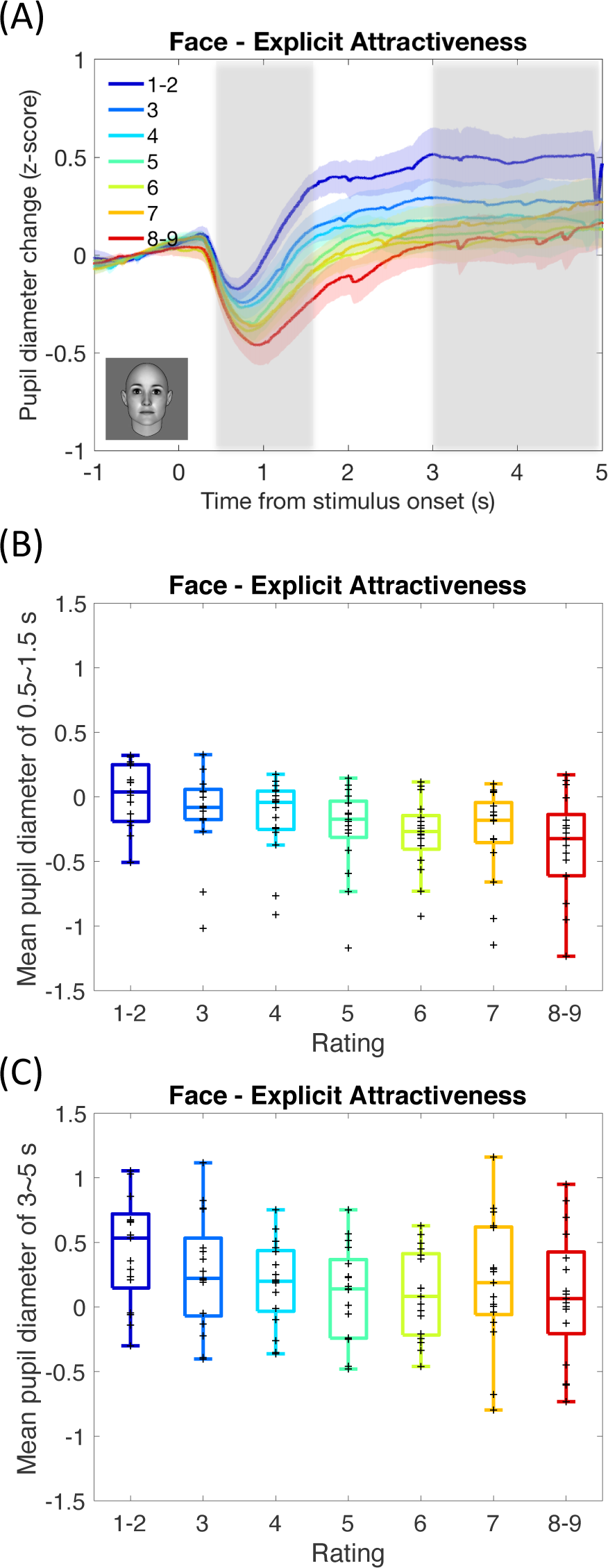
Pupil response results in Experiment 3 (delayed response). (A) Mean pupil diameter change as a function of time reference to the target onset during the delayed attractiveness judgment for faces (as shown in a sample image). Curves are parameterized with rating score depending on individual participants’ choices (1 for least attractive and 9 for most attractive). The color shadows represent standard errors among participants. The gray shadow represents the time window for averaging the pupil size to present the amount of pupil response for statistical analysis (see Materials and Methods for details and Table 3 for results). (B)–(C) Box plots of the mean pupil size over the specified time window (0.5–1.5 s for panel B and 3–5 s for panel C). The plus signs represent individual data.

**Figure 4.**
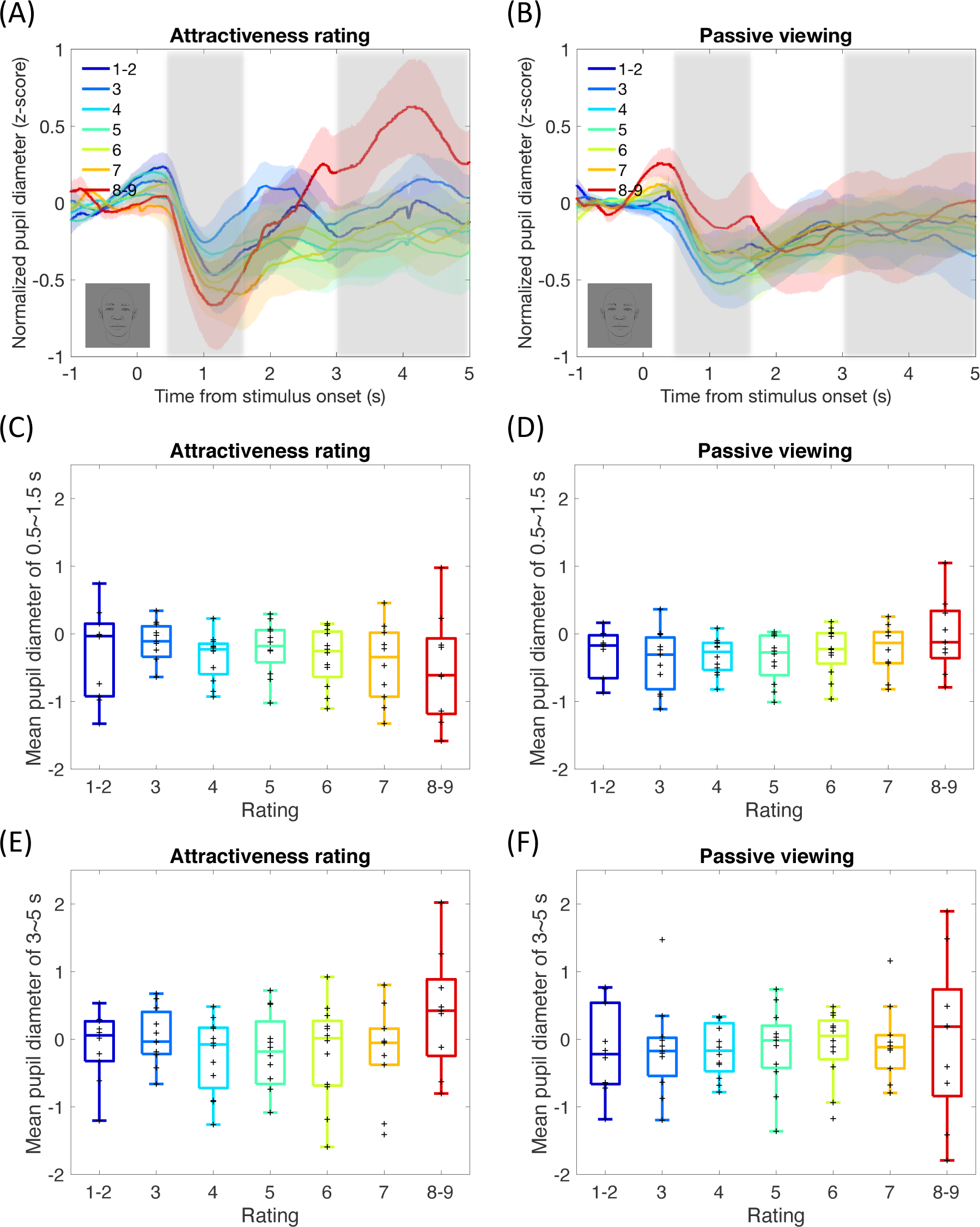
Pupil response results in Experiment 4 (delayed response or passive viewing with line-drawing faces, as shown in sample images). (A)–(B) Mean pupil diameter change as a function of time reference to the target onset during the delayed attractiveness judgment for faces (panel A) or passive viewing (panel B). Curves are parameterized with rating score depending on individual participants’ choices (1 for least attractive and 9 for most attractive). The color shadows represent standard errors among participants. The gray shadow represents the time window for averaging the pupil size to present the amount of pupil response for statistical analysis (see Materials and Methods for details and Table 3 for results). (C)–(F) Box plots of the mean pupil size over the specified time window in the specified task demand conditions (0.5–1.5 s in the attractiveness rating condition for panel C, 0.5–1.5 s in the passive viewing condition for panel D, 3–5 s in the attractiveness rating condition for panel E, and 3–5 s in passive viewing condition for panel F). The plus signs represent individual data.

**Figure 5.**
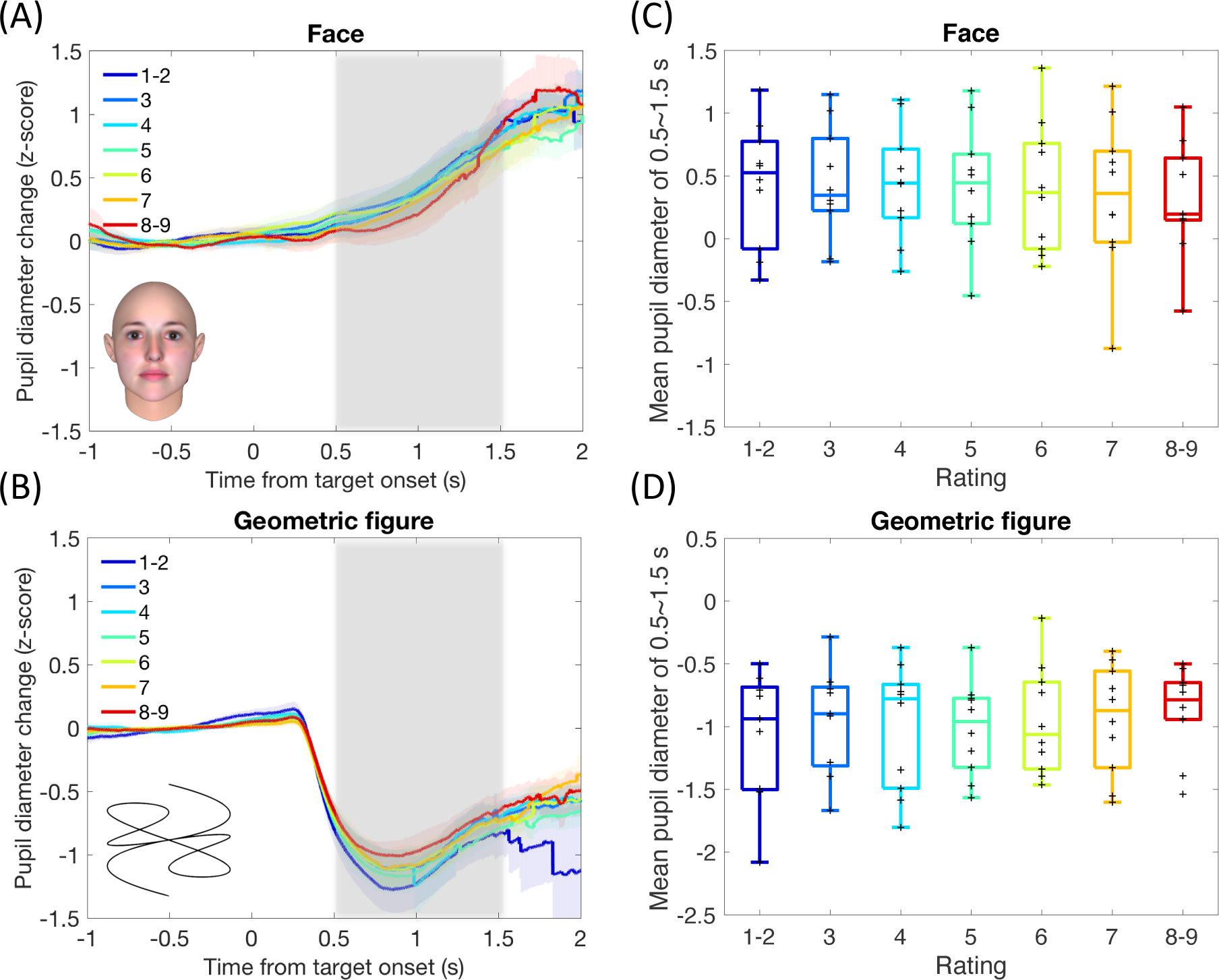
Pupil response results in Experiment 5 (overall pupil dilation to faces and overall pupil constriction to geometric figures). (A)–(B) Mean pupil diameter change as a function of time reference to the target onset during attractiveness judgment for faces (panel A) or geometric figures (panel B). Curves are parameterized with rating score depending on individual participants’ choices (1 for least attractive and 9 for most attractive). The color shadows represent standard errors among participants. The gray shadow represents the time window for averaging the pupil size to present the amount of pupil response for statistical analysis (see Materials and Methods for details and Table 3 for results). (C)–(D) Box plots of the mean pupil size over the specified time window in the face condition (panel C) and geometric figure condition (panel D). The plus signs represent individual data.

**Figure 6.**
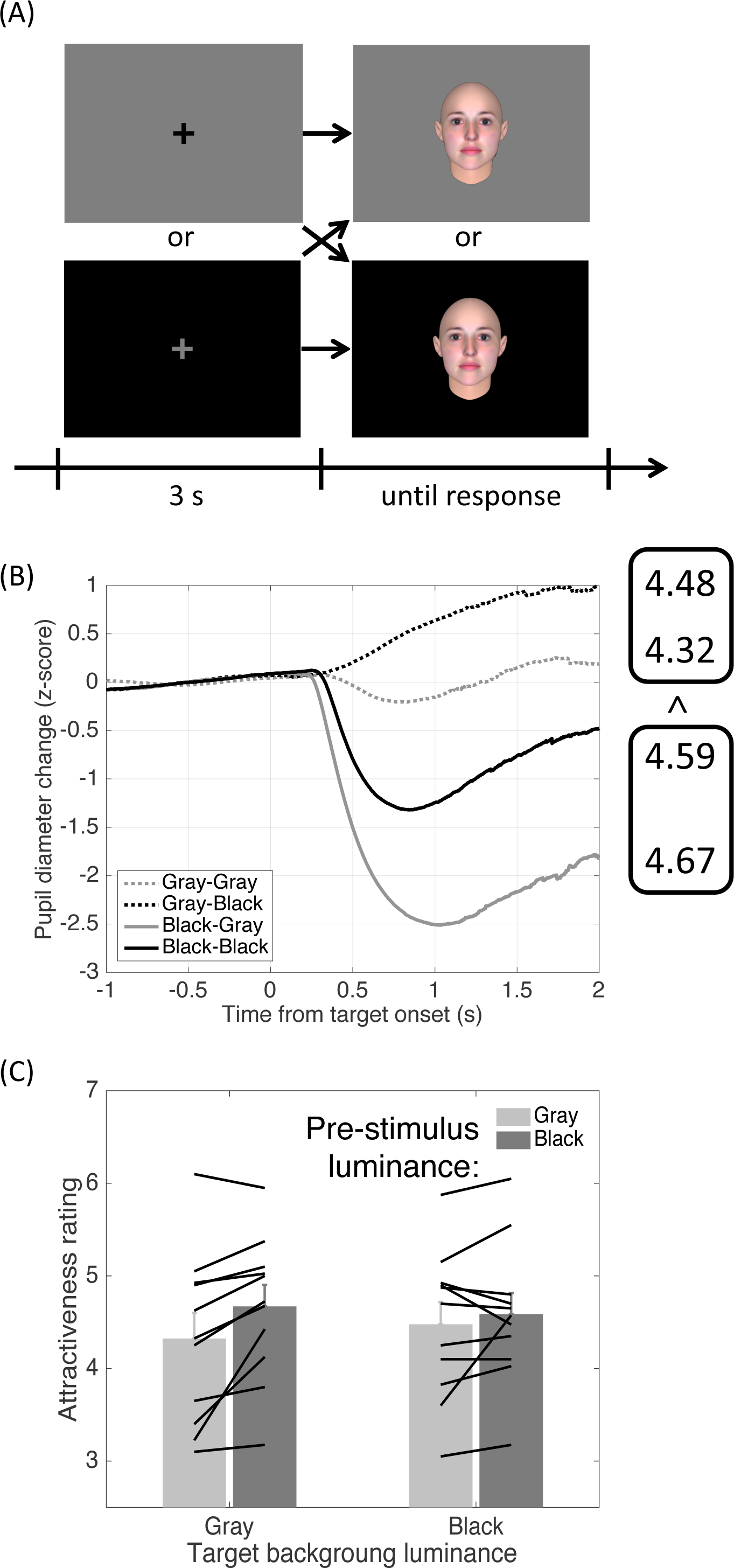
Procedure and results in Experiment 6. (A) Illustration of experimental procedure (not to scale). (B) Pupil response results: mean pupil diameter change as a function of time reference to the target onset during the facial attractiveness judgment. Curves are parameterized with pre-stimulus and target background luminance conditions. Dotted lines represent the gray pre-stimulus condition. Solid lines represent the black pre-stimulus conditions. Gray lines represent the gray target background conditions. Black lines represent the black target background conditions. The numbers on the right are mean attractiveness rating scores corresponding to the pre-stimulus and target background luminance conditions. (C) Mean attractiveness rating as a function of pre-stimulus and target background luminance. Each black line represents individual participants’ mean rating score. Error bars represent standard errors among participants.

In Experiment 7, we examined whether sequential luminance contrast alone, when not inducing a strong difference in pupil response, causes differences in attractiveness judgments. We divided visual fields into two halves (left/right) with luminance disparities in the fixation display and then presented the target image to the left or right visual field (see Figure 7A). In this case, there was sequential luminance contrast to the target image (different luminance conditions depending on the spatial relationship between the target image location and the fixation display’s luminance disparities, i.e., black on the left or the right visual fields), but the overall average luminance of the fixation display remained the same to induce a similar pupillary light reflex (to the face display with a gray background). Participants were allowed to move their gaze to the face position (Experiment 7a) or were instructed to always fixate the center even when the face was presented peripherally (Experiment 7b).

**Figure 7.**
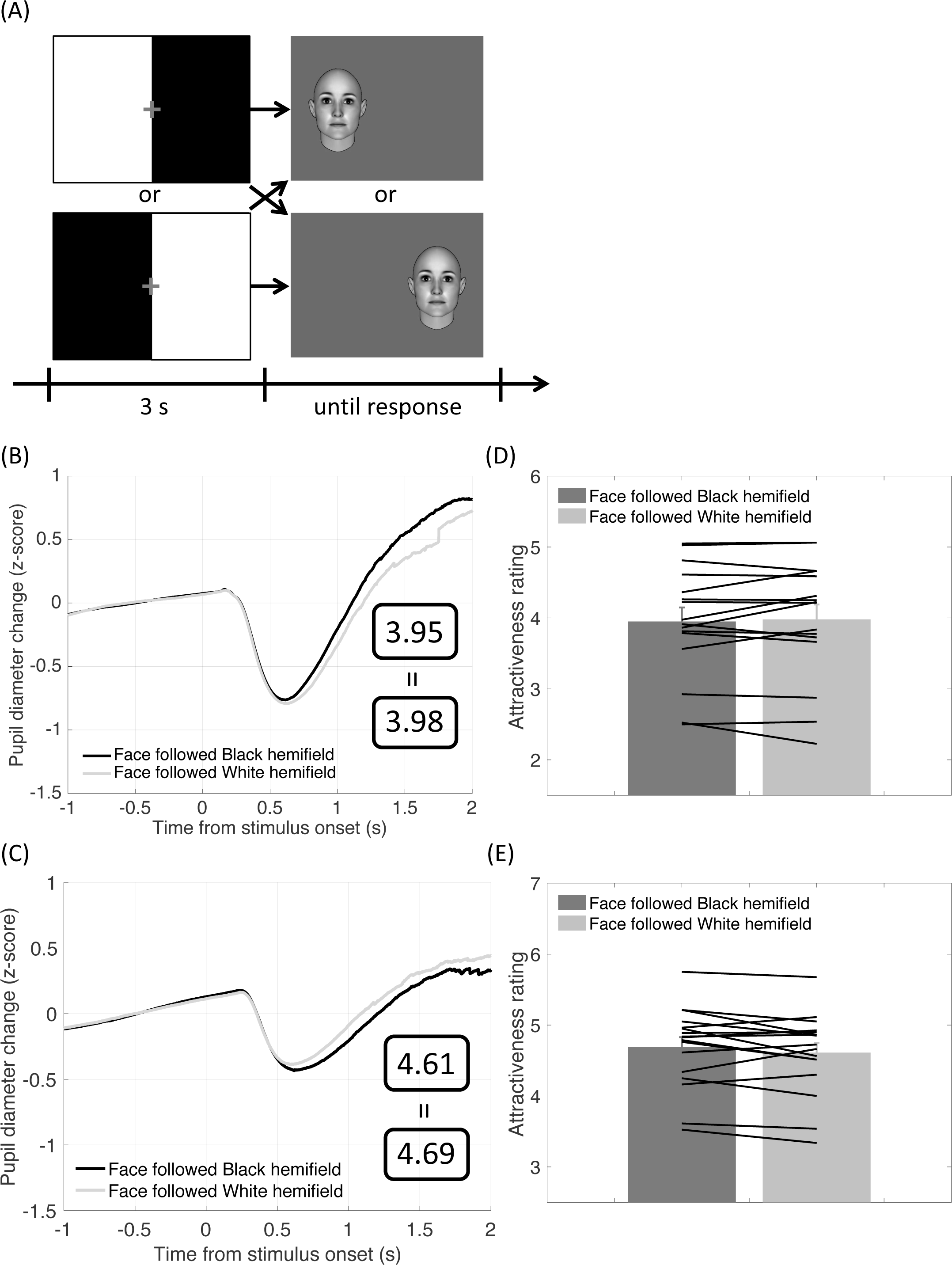
Procedure and results in Experiment 7. (A) Illustration of experimental procedure (not to scale). (B)–(C) Pupil response results in Experiment 7a (panel B) and 7b (panel C): mean pupil diameter change as a function of time reference to the target onset during the facial attractiveness judgment. Curves are parameterized with the relationship between the target face and pre-stimulus hemifield’s luminance conditions. The numbers in the squares are the mean attractiveness rating scores corresponding to the pre-stimulus hemifield’s luminance conditions. (D)-(E) Mean attractiveness rating as a function of pre-stimulus’s hemifield’s luminance in Experiment 7 when (panel D) eye movement was allowed or (panel E) eye movement was not allowed (participants fixated the central fixation cross throughout the trial). Each black line represents individual participants’ mean rating score. Error bars represent standard errors among participants.

### Participants

Sixty-eight adults (43 females, age range of 20–48, median age = 35 years) participated in the current study: 13 in Experiment 1, 15 in Experiment 2, 17 in Experiment 3, 12 in Experiment 4, 10 in Experiment 5, 11 in Experiment 6 (the same group of participants as in Experiment 1 with two excluded due to the data lost by program error for the first two participants), 16 in Experiment 7a (the same group of participants as in Experiment 2, plus one who was excluded from Experiment 2 due to data recording loss for the last participant) and 17 in Experiment 7b (the same group of participants as in Experiment 3). All had normal or corrected-to-normal vision and were naïve about the purpose of the experiments. The current study was performed in accordance with the Declaration of Helsinki. All participants gave written informed consent before the experiment and received payment for their participation.

### Apparatus and stimuli

Visual stimuli were presented on an 18.1-inch monitor (Eizo FlexScan L685Ex) with a 60-Hz frame rate, controlled by a personal computer (Dell OptiPlex 755). In Experiment 1, in the target display, a target image was presented at the center of the screen against a gray background (21.04 cd/m^2^). There were three conditions. In the face-attractiveness and face-roundness conditions, the target image was a face (6.42° width × 7.83° height), which was generated by FaceGen (Singular Inversions Inc.) software. Faces consisted of eight subcategories of the combination of two races (Asian or European), gender, and age range (old or young). There were 20 face images in each subcategory, thus 160 face images in total (mean luminance = 25.57 cd/m^2^: maximum of 43.13 cd/m^2^; minimum of 12.82 cd/m^2^). In the geometric figure-attractiveness condition, a geometric figure, in black lines, with 10.62° width × 7.83° height was presented at the center of the screen to serve as the target. The figures were Fourier descriptors generated by a Matlab program (MathWorks Inc.) with properties specified and varied as a combination of symmetry (symmetric or asymmetric) and simplicity (simple or complex). A total of 160 geometric figures were generated (mean luminance = 16.46 cd/m^2^: maximum of 23.22 cd/m^2^; minimum of 4.27 cd/m^2^). All the target displays were interlaid with a fixation display, which consisted of a black fixation cross (0.5° × 0.5°, 0.35 cd/m^2^) against a gray background.

In Experiment 2, the stimuli and experimental structure were the same as in Experiment 1, except that instead of the geometric figures, we used natural scene images. The original images were color photos collected from public websites. They consisted of eight subcategories: animal, food, flower, mountain, sky, lake, ocean, and desert. There were 20 images in each subcategory, thus 160 images in total. The size of the images was within 8° width or 9.75° height, presented at the center of the screen. The original color natural scenes and face images (used in Experiment 1) were modified to be in the gray scale with the same mean luminance as the background (21.04 cd/m^2^) by using the SHINE toolbox (Willenbockel et al., 2010).

In Experiment 3, the same set of face images as in Experiment 2 was used. No natural scene images were used in Experiment 3. In Experiment 4, forty face images (five in each subcategory as used in Experiment 1) were selected for further processing. Each face was manually line-drawn based on the original image with the pupil adjusted in five different sizes (see example in Figure 8). In Experiment 5, the stimuli were the same as in Experiment 1, except for the following modifications. First, the faces were presented at the center of the screen against a white background (94.04 cd/m^2^). The interlaid fixation display for faces consisted of a light gray fixation cross (58.66 cd/m^2^) against the white background. Second, for geometric figures, the background was black (0.35 cd/m^2^), and the fixation cross was dark gray (2.44 cd/m^2^) in the interlaid fixation display.

**Figure 8.**
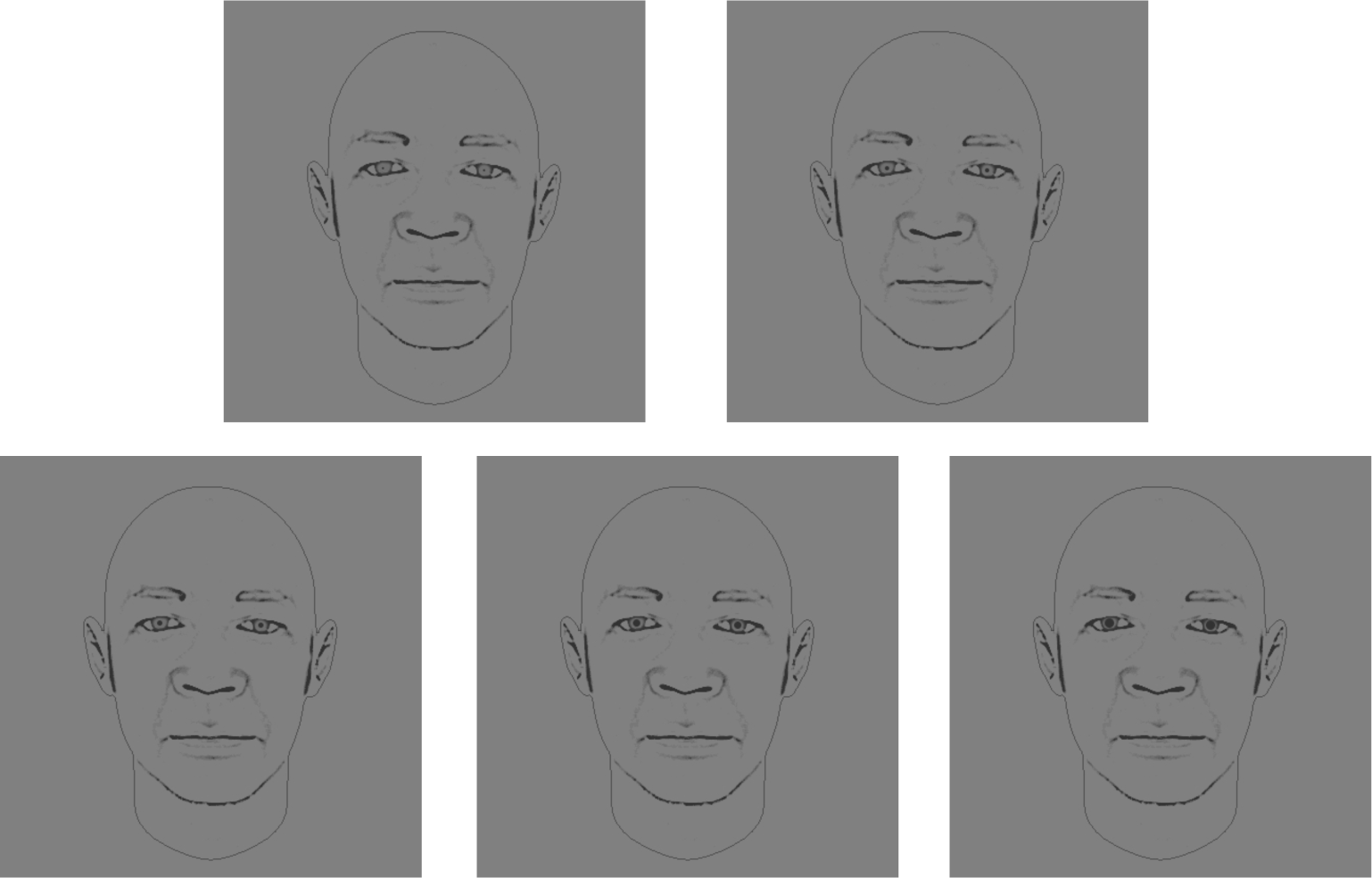
Sample line-drawing faces used in Experiment 4. The five different faces have five different pupil sizes.

In Experiment 6, we used the faces that were judged as median attractive by individual participants in Experiment 1. For each participant and in each race subcategory (combined across gender and age), the rating scores were ranked order, and the 20 faces that corresponded to the median attractive rank order were used. No geometric figures were used in Experiment 6. Faces were presented at the center of the screen against a gray (21.04 cd/m^2^) or a black (0.35 cd/m^2^) background. The interlaid fixation display consisted of a black fixation cross (0.5° × 0.5°, 0.35 cd/m^2^) against the gray background or a gray fixation cross (0.5° × 0.5°, 21.04 cd/m^2^) against the black background.

In Experiment 7a, following the same procedure as in Experiment 6, for each participant we selected 40 median attractive faces (20 for each race) used in Experiment 2 based on individual judgments. In Experiment 7b, the faces were selected based on the individual judgments in Experiment 3. In both Experiment 7a and 7b, no natural scene images were used. Faces were presented to the left or right visual field with 5.03° of eccentricity against the gray background (21.04 cd/m^2^). In Experiment 7a, the interlaid fixation display consisted of a gray fixation cross (21.04 cd/m^2^) against the background with luminance disparity across the visual field: black (0.35 cd/m^2^) on the left and white (94.04 cd/m^2^) on the right or vice versa. In Experiment 7b, the fixation cross was red and remained visible during the face target presentation.

### Design

In all experiments, each trial consisted of the target display following the fixation display presented for 3 s. In Experiments 1 and 2, the three conditions were conducted as within-subject factors in different blocks with counterbalanced order among the participants. In the face-attractiveness and face-roundness conditions, the faces of different races (Asian and European) where presented in separated subblocks. Each subblock consisted of 80 face images presented in randomly assigned order. There was no break between the subblocks. In the geometric figure-attractiveness condition (in Experiment 1), all 160 geometric figures were presented in randomly assigned order. In the natural scene-attractiveness condition (in Experiment 2), the images of different subcategories were presented in separate subblocks without a break between them. The order of the images within subblocks and the order of the subcategories were randomized.

In Experiment 3, only the face-attractiveness condition was conducted. The face was presented on the screen for 5 s and then replaced by the fixation display. In Experiment 4, we applied the nested design in which each participant viewed 40 face images only, all with different identities. Each pupil size level was presented eight times with different face identities. There were two conditions: attractiveness rating and passive viewing. The two conditions were conducted as within-subject factors in separate blocks with the order counterbalanced across the participant. The same set of stimuli was used in the two conditions, with the trial order randomly assigned. The face image was presented with its luminance contrast gradually increased for 1 s, stayed at the center of the screen for 3 s, and then gradually disappeared for 1 s. In Experiment 5, the design was the same as in Experiment 1 except that there were only two conditions: face-attractiveness and geometric figure-attractiveness conditions (no face-roundness condition).

In Experiments 6, 7a, and 7b, the two types of fixation display and the two types of target display were conducted as within-subject factors. In each race subcategory, the 20 median attractive faces were presented for four times, in each of the 2 (fixation display) × 2 (target display) conditions. There were thus 80 trials in each race subcategory and 160 trials in total. As in Experiment 1, the faces of different races were presented in different subblocks in randomly assigned order. In each subblock, the 80 trials with different manipulation conditions were presented in randomly assigned order.

### Procedure

Participants sat in front of the monitor at an 80-cm distance with their head supported on a chin-rest. In each session/experimental block, participants went through the five-point Eyelink calibration program to calibrate and validate their eye data. After the calibration procedure, the experiment started without practice trials. Participants were instructed to fixate the central fixation cross during the fixation display. Once the target display was shown, they were asked to make a judgment (attractiveness or roundness) on the target image (faces, geometric figures, or natural scenes). In Experiments 1, 2, 5, 6, 7a, and 7b, they were free to take their own pace in making the decision. In Experiments 3 and 4, they were asked to give the rating score after the face disappeared. In the passive viewing condition of Experiment 4, they were asked to look at the faces without any task involved. In all experiments except Experiment 7b, they were allowed to move their gaze to the stimulus position. In Experiment 7b, they were instructed to fixate the central fixation cross even when the face was presented peripherally. In the attractiveness judgment condition, participants rated how attractive the face, geometric figure, or natural scene was, i.e., how much they liked the given image. In the roundness judgment condition, participants rated how round the face was. They indicated their answer by pressing the number pad on a keyboard from 1 (least attractive/round) to 9 (most attractive/round). They were encouraged to use all nine numbers if possible but not necessarily equate the distribution so that they would make their judgment naturally. After they gave their answer or 0.5 s after the stimulus presentation in the passive condition of Experiment 4, the next trial started with the 3-s fixation display. Participants made judgments for all trials straight without break. Each session took about 20 minutes, and there was more than a 20-minute break between the sessions.

### Eye metrics data analysis

Eye movements (including pupillary responses and gaze location) were recorded binocularly with an infrared eye-tracker camera (Eyelink 1000 Desktop Mount, SR Research Ltd.). The camera was positioned below the monitor. The sampling rate of the recording was 1000 Hz. Since pupillary responses are consensual, only data from the right eye were used. Data during blinks was interpolated using shape-preserving piecewise cubic function. During the time window of -1 to 2-s reference to stimulus onset, blinks accounted for 15.1% of data points in Experiment 1, 8.4% in Experiment 2, 19.1% in Experiment 5, 18.3% in Experiment 6, 11.6% in Experiment 7a, and 16.5% in Experiment 7b. The blink rate during the -1 to 5-s time window reference to the target onset was 11.7% and 21.4% for Experiments 3 and 4, respectively. The blink rate was in the normal range when natural blinking was allowed (e.g., Goldinger and Papesh, 2012). To compare the pupillary response results across participants and conditions, pupil diameter data were normalized using all the data recorded in each session and baseline corrected by subtracting the mean of the data during the 1-s period before the stimulus onset. For the gaze-contingent pupillary response analysis, in each trial, all the gazes, i.e., the fixations that were detected from the Eyelink system, were first identified (mean duration = 421 ms in Experiments 1-5). The normalized pupil diameter data were then baseline corrected by subtracting the mean of the data during the 100-ms period before the gaze onset. For the gaze location analysis, the gaze location data were averaged during target presentation, i.e., 0–2 s after the target onset in Experiments 1, 2, 5, and 7, and 0–5 s in Experiments 3 and 4. The gazed local luminance was calculated by averaging the luminance of the image regions within 1 degree of visual angle of the gazing point across time.

### Statistical analysis

#### Experiments 1–5

For behavioral responses, mean reaction times for all experiments (except for Experiments 3 and 4 since the response was held and postponed) are shown in Table 2. On average, participants made decisions at around 2 s. The histograms of rating for Experiments 1 and 2 are shown in Figures 9 and 10, respectively. In all the conditions we tested, extreme rating scores (i.e., 1 and 9) were least given, with only about half the frequency of their adjacent scores (i.e., 2 and 8, respectively). We therefore combined trials with rating scores of 1 and 2 and those with scores of 8 and 9 together to reduce the noise due to few trials and to balance the trial numbers across conditions for further analysis.

**Figure 9.**
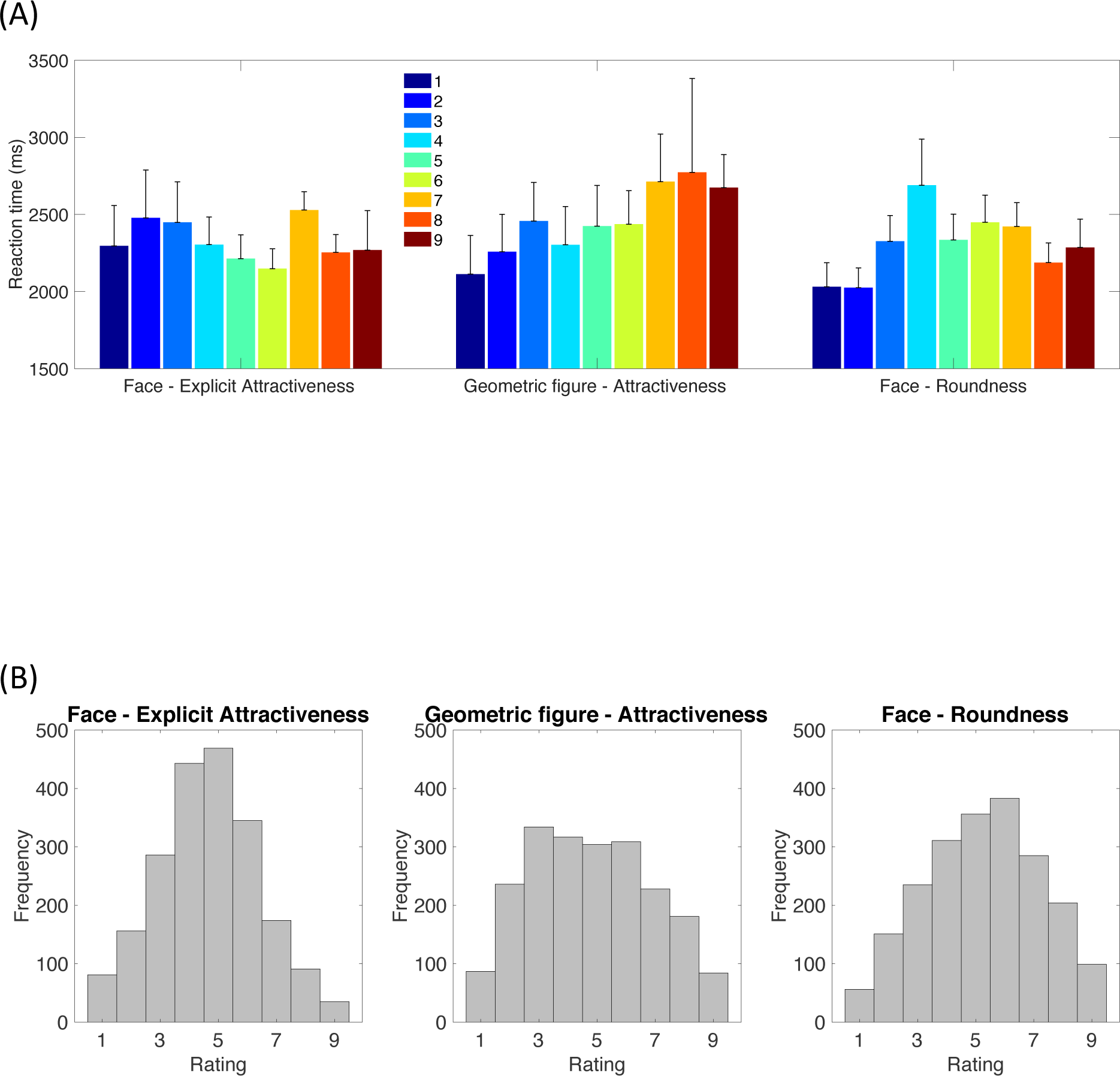
Behavioral results in Experiment 1. (A) Mean reaction times as a function of rating scores. Error bars represent standard errors among participants. (B) Histograms of rating scores.

**Figure 10.**
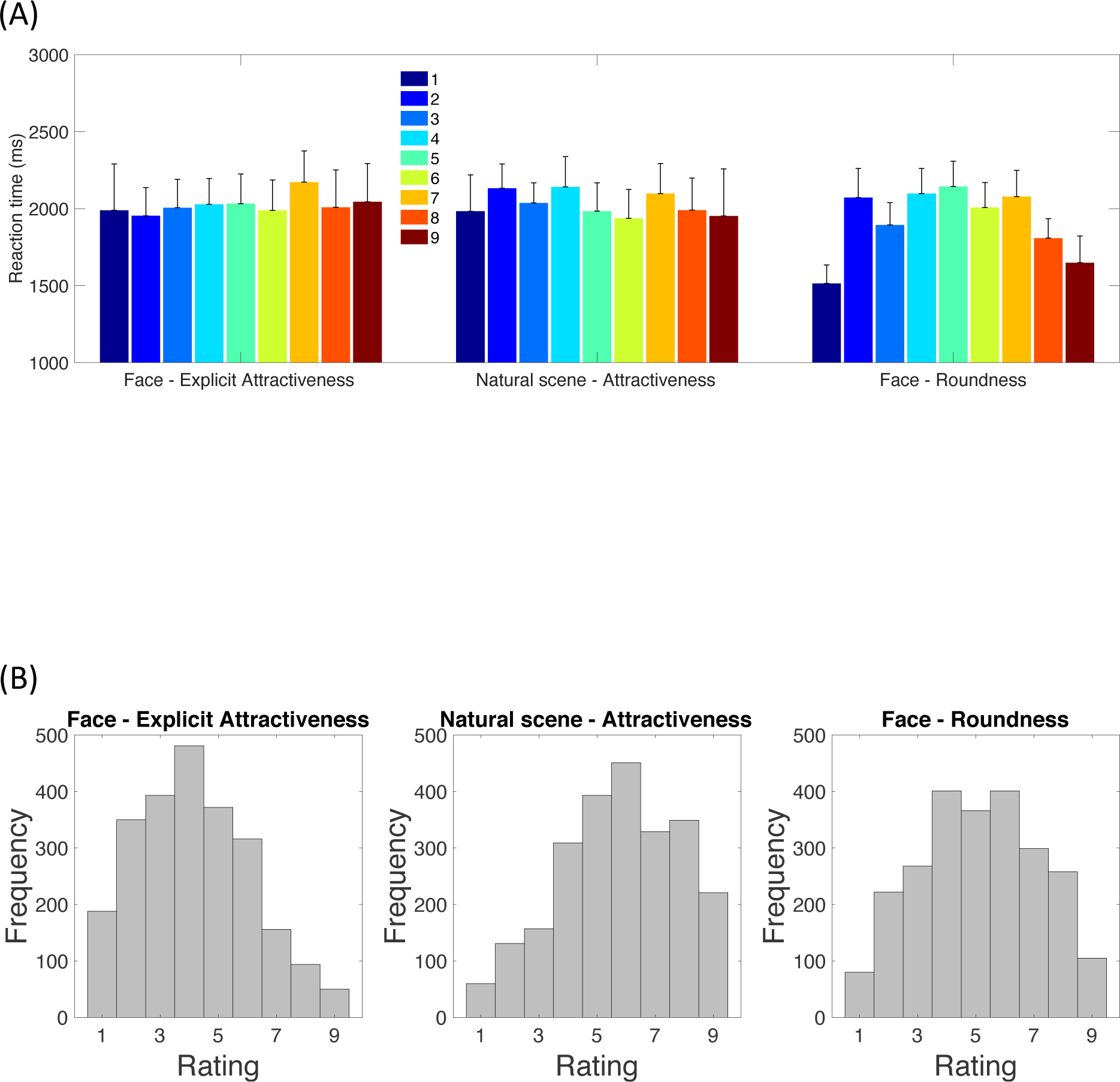
Behavioral results in Experiment 2. (A) Mean reaction times as a function of rating scores. Error bars represent standard errors among participants. (B) Histograms of rating scores.

**Table 2.**
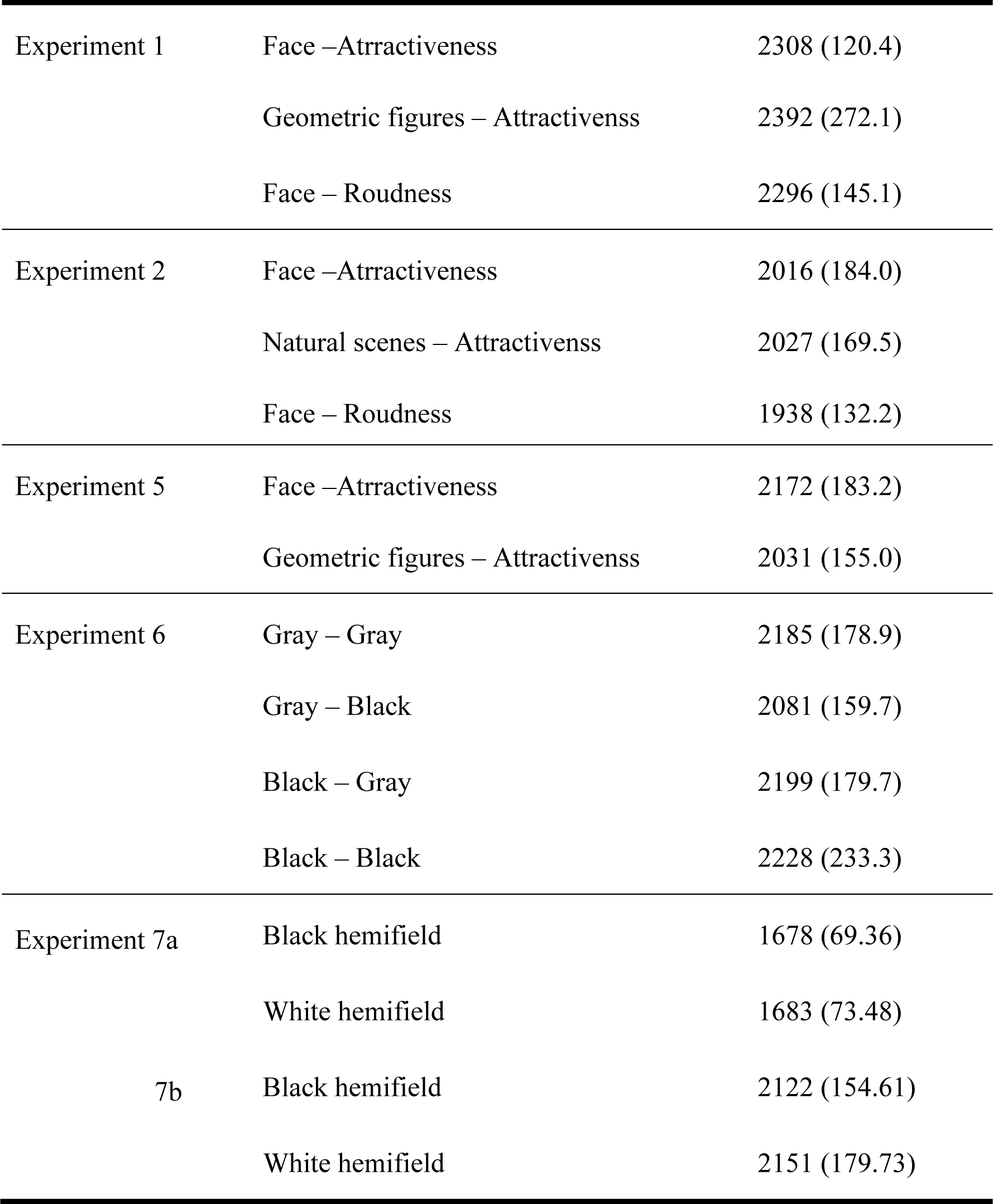
Mean reaction times (ms) under each condition in all experiments except Experiments 3 and 4. Numbers in parentheses are standard errors among participants (also see Figures 9 and 10).

We averaged the pupil diameter 0.5–1.5 s or 3–5 s (only for Experiments 3 and 4) after stimulus onset to represent the pupil constriction/dilation response. Mean pupil diameter data was subjected to a repeated-measure ANOVA with the 7-level rating score as the within-subject factor in each condition. We also examined the correlation between the mean pupil diameter and the attractiveness rating on a trial-by-trial basis. Pearson’s correlation coefficients between the mean pupil diameter and rating score for individual participants were calculated and subjected to a one-sample t-test to examine if they deviated from zero. Partial correlation coefficients between the mean pupil diameter and rating score were also calculated when the effect of mean gazed local luminance over the same time window as the mean pupil diameter was removed. Results of the ANVOA, linear trend analysis, and correlation analyses (with and without the mean gazed local luminance being partial out) are shown in Table 3.

**Table 3.**
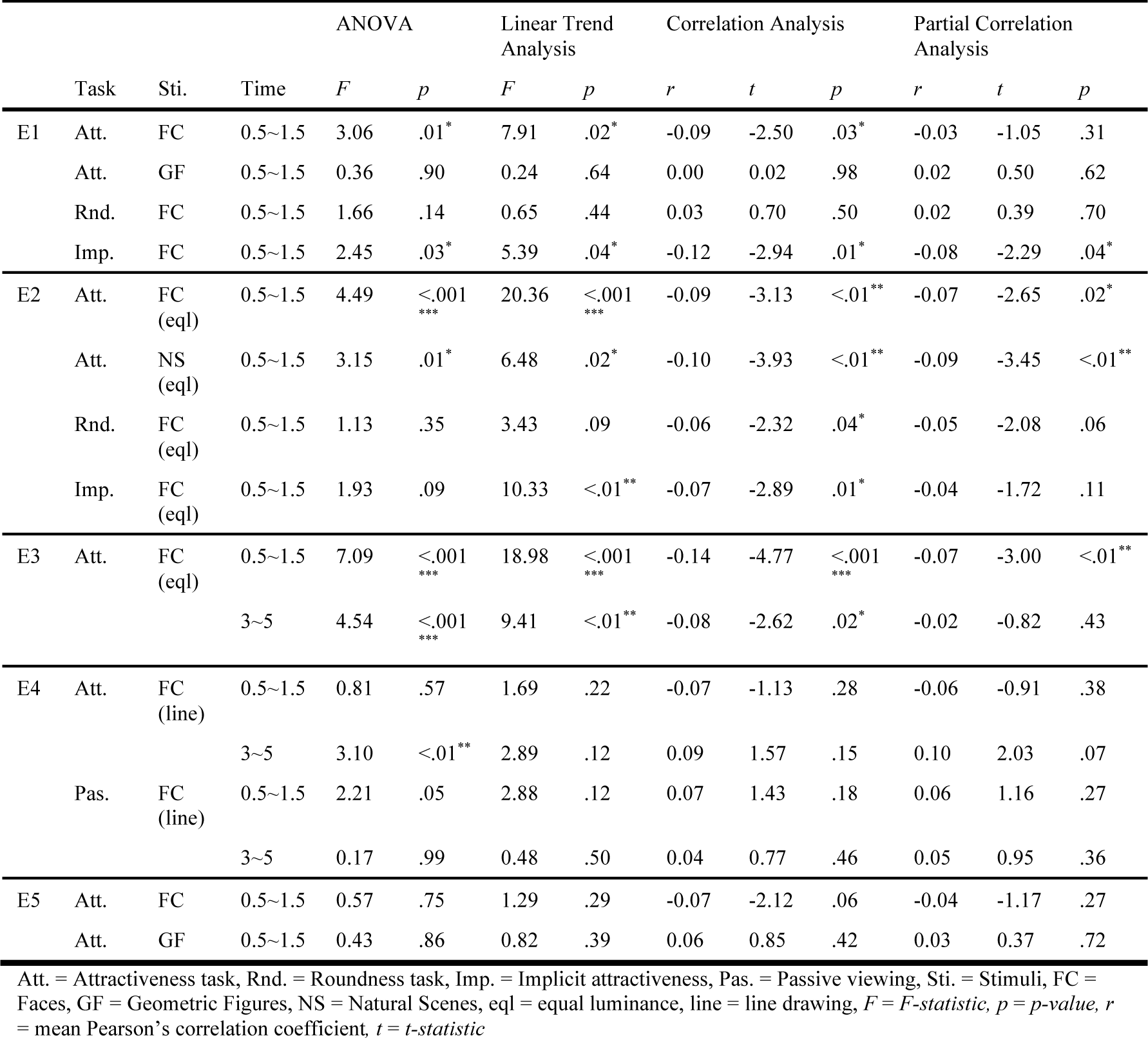
Statistical results for the mean pupil size during specified time window (s) in Experiments 1-5. ANOVA examined whether pupil size differed among ratings. Linear Trend Analysis examined whether pupil size linearly correlated with rating using average data. Correlation Analysis and Partial Correlation Analysis examined the correlation between pupil size and rating on a trial-by-trial base, without and with the effect of gazed local luminance being removed, respectively.

|  | Task | Sti. | Time | ANOVA |  | Linear Trend Analysis |  | Correlation Analysis |  |  | Partial Correlation Analysis |  |  |
| --- | --- | --- | --- | --- | --- | --- | --- | --- | --- | --- | --- | --- | --- |
|  |  |  |  | *F* | *p* | *F* | *p* | *r* | *t* | *p* | *r* | *t* | *p* |
| E1 | Att. | FC | 0.5~1.5 | 3.06 | .01* | 7.91 | .02* | -0.09 | -2.50 | .03* | -0.03 | -1.05 | .31 |
|  | Att. | GF | 0.5~1.5 | 0.36 | .90 | 0.24 | .64 | 0.00 | 0.02 | .98 | 0.02 | 0.50 | .62 |
|  | Rnd. | FC | 0.5~1.5 | 1.66 | .14 | 0.65 | .44 | 0.03 | 0.70 | .50 | 0.02 | 0.39 | .70 |
|  | Imp. | FC | 0.5~1.5 | 2.45 | .03* | 5.39 | .04* | -0.12 | -2.94 | .01* | -0.08 | -2.29 | .04* |
| E2 | Att. | FC (eq) | 0.5~1.5 | 4.49 | <.001*** | 20.36 | <.001*** | -0.09 | -3.13 | <.01** | -0.07 | -2.65 | .02* |
|  | Att. | NS (eq) | 0.5~1.5 | 3.15 | .01* | 6.48 | .02* | -0.10 | -3.93 | <.01** | -0.09 | -3.45 | <.01** |
|  | Rnd. | FC (eq) | 0.5~1.5 | 1.13 | .35 | 3.43 | .09 | -0.06 | -2.32 | .04* | -0.05 | -2.08 | .06 |
|  | Imp. | FC (eq) | 0.5~1.5 | 1.93 | .09 | 10.33 | <.01** | -0.07 | -2.89 | .01* | -0.04 | -1.72 | .11 |
| E3 | Att. | FC (eq) | 0.5~1.5 | 7.09 | <.001*** | 18.98 | <.001*** | -0.14 | -4.77 | <.001*** | -0.07 | -3.00 | <.01** |
|  |  |  | 3~5 | 4.54 | <.001*** | 9.41 | <.01** | -0.08 | -2.62 | .02* | -0.02 | -0.82 | .43 |
| E4 | Att. | FC (line) | 0.5~1.5 | 0.81 | .57 | 1.69 | .22 | -0.07 | -1.13 | .28 | -0.06 | -0.91 | .38 |
|  |  |  | 3~5 | 3.10 | <.01** | 2.89 | .12 | 0.09 | 1.57 | .15 | 0.10 | 2.03 | .07 |
|  | Pas. | FC (line) | 0.5~1.5 | 2.21 | .05 | 2.88 | .12 | 0.07 | 1.43 | .18 | 0.06 | 1.16 | .27 |
|  |  |  | 3~5 | 0.17 | .99 | 0.48 | .50 | 0.04 | 0.77 | .46 | 0.05 | 0.95 | .36 |
| E5 | Att. | FC | 0.5~1.5 | 0.57 | .75 | 1.29 | .29 | -0.07 | -2.12 | .06 | -0.04 | -1.17 | .27 |
|  | Att. | GF | 0.5~1.5 | 0.43 | .86 | 0.82 | .39 | 0.06 | 0.85 | .42 | 0.03 | 0.37 | .72 |

Att. = Attractiveness task, Rnd. = Roundness task, Imp. = Implicit attractiveness, Pas. = Passive viewing, Sti. = Stimuli, FC = Faces, GF = Geometric Figures, NS = Natural Scenes, eq = equal luminance, line = line drawing, *F* = *F*-statistic, *p* = *p*-value, *r* = mean Pearson's correlation coefficient, *t* = *t*-statistic

#### Effect of task order between attractiveness and roundness tasks on pupil constriction under explicit- and implicit-attractiveness conditions (Experiments 1 and 2 only)

Mean pupil diameter data sorted by facial attractiveness was subjected to a repeated-measure ANOVA with the 7-level rating score as the within-subject factor and task order (attractiveness judgment first or roundness judgment first) as the between-subject factor. In Experiment 1, in the Explicit-Attractiveness condition, the effect of rating was significant [*F*(6,66) = 2.99, *p* < .02], but not the effect of task order [*F*(1,11) = 1.38, *p* = .26] or the interaction between rating and task order [*F*(6,66) = 0.74, *p* = .62]. The linear trend of the rating was significant [*F*(1,11) = 7.48, *p* < .02] and did not interact with task order [*F*(1,11) = 0.34, *p* = .57]. The same pattern of results was found in the Implicit-Attractiveness condition: the effect of rating was significant [*F*(6,66) = 2.49, *p* < .04], but not the effect of task order [*F*(1,11) = 1.52, *p* = .24] or the interaction [*F*(6,66) = 1.21, *p* = .31]. In contrast, the linear trend or rating was significant [*F*(1,11) = 7.06, *p* < .03], with a marginally significant interaction with the task order [*F*(1,11) = 4.73, *p* = .05]. The effect of the linear trend was significant when the participant performed the roundness task earlier than the attractiveness task [*F*(1,5) = 8.42, *p* < .04] but not the other way around [*F*(1,6) = 0.93, *p* = .37].

In Experiment 2, in the Explicit-Attractiveness condition, the effect of rating was significant [*F*(6,78) = 4.72, *p* < .001] but not the effect of task order [*F*(1,13) = 2.61, *p* = .13] or the interaction [*F*(6,78) = 1.70, *p* = .13]. The linear trend of rating was significant [*F*(1,13) = 18.54, *p* < .001] and did not interact with task order [*F*(1,13) = 0.25, *p* = .62]. In the Implicit-Attractiveness condition, none of the main effects [*F*(6,78) = 1.93, *p* = .09 and *F*(1,13) = 0.82, *p* = .38 for the effect of rating and task order, respectively] or interaction were significant [*F*(6,78) = 1.01, *p* = .42]. The linear trend of rating was significant [*F*(1,13) = 9.54, *p* < .01] and did not interact with task order [*F*(1,13) = 0.08, *p* = .79].

#### Experiment 6

Mean pupil diameter data (i.e., average pupil diameter data 0.5–1.5 s after the stimulus onset) was subjected to a two-way ANOVA with pre-stimulus luminance (i.e., the fixation display) and target background luminance as within-subject factors. The same ANOVA test was conducted using the mean facial attractiveness rating scores. Partial correlation analyses were conducted among the following three variables: pre-stimulus pupil baseline, pupil constriction amplitude, and attractiveness rating. Pre-stimulus pupil baseline was calculated by averaging pupil diameter data 0–1 s before the stimulus onset. Pupil constriction amplitude, i.e., the amount of pupil constriction change responding to the stimulus, was calculated by subtracting the pre-stimulus pupil baseline from the mean pupil diameter data 0.5–1.5 s after the stimulus onset. Linear partial correlation coefficients for individual participants were calculated and subjected to a one-sample t-test to examine if they deviated from zero.

#### Experiment 7

Mean pupil diameter data (i.e., average pupil diameter data 0.5–1.5 s after the stimulus onset) in the two conditions where the face followed white and black hemifield, respectively, were subjected to a paired two-sample t-test. The same statistical test was conducted using the mean facial attractiveness rating scores.

## Results

### Transient pupil constriction reflects attractiveness

In Experiment 1, the pupil in general constricted in response to the presentation of the faces. Most importantly, during the inspection, the degree of pupil constriction in each participant was linearly correlated with the facial attractiveness rated immediately after the inspection in every single trial [see Figure 1A for overall pupillary response change and Figure 1E for box plot for statistical analysis: *F*(1,12) = 7.91, *p* < .03 for linear trend analysis]. The more attractive the face was, the more the pupils constricted. In contrast, the amount of pupil constriction did not correlate with attractiveness judgments for geometric figures [*F*(1,12) = 0.24, *p* = .64; see Figure 1B and 1F] or roundness judgments for faces [*F*(1,12) = 0.65, *p* = .44; see Figure 1C and 1G]. Intriguingly, when the faces were sorted by their attractiveness (although the explicit task demand was to judge their roundness), the degree of pupil constriction showed a linear correlation with the implicit, or task-irrelevant, attractiveness of the faces [*F*(1,12) = 5.39, *p* < .05; see Figure 1D and 1H]—the same pattern of results as when the task demand was to judge the attractiveness (i.e., the face-attractiveness condition). Note again that the order of the three conditions (face-attractiveness, geometric figure-attractiveness, and face-roundness) was counterbalanced across participants. A further analysis involving condition order as a factor (see Material and Methods for details) showed that the pattern of the pupillary responses to facial attractiveness (either explicit or implicit) remained the same regardless of the condition order (*p*s > .05); it did not matter whether the faces were judged on attractiveness earlier than roundness or vice versa. In summary, the overall results of Experiment 1 suggest that the pupil constriction response to facial attractiveness is task-specific (in contrast to roundness judgment), automatic, and free of memory. A potential problem in Experiment 1, however, is that we controlled luminance across stimuli rather crudely (see sample images in Figure 11), and it may be criticized that the result could be explained by the low-level factor since the pupil is very sensitive to luminance (contrast).

**Figure 11.**
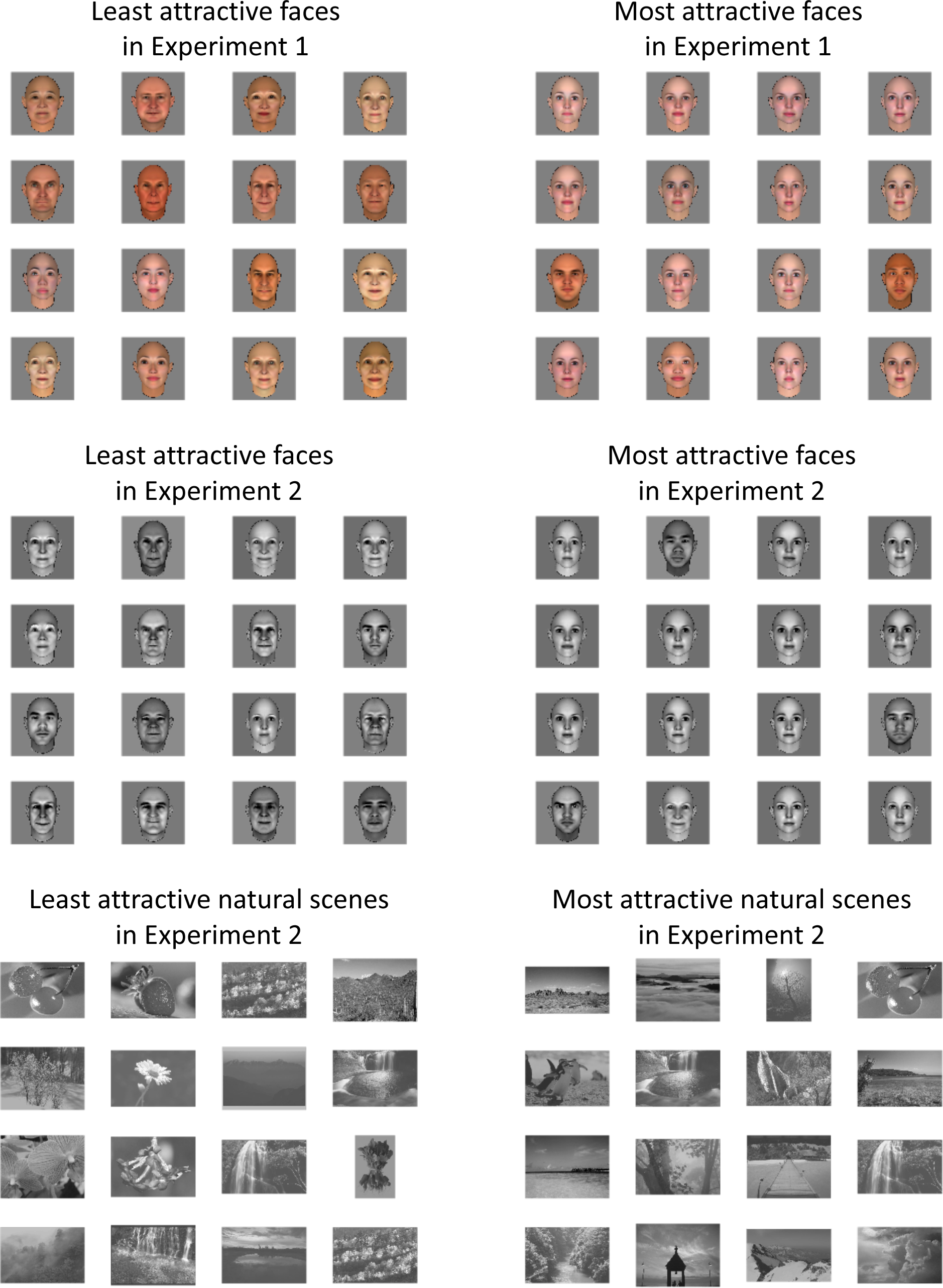
Sample images which were rated as least attractive (rating score = 1) and most attractive (rating score = 9) in Experiments 1 and 2. As shown in the examples, the same image (e.g., the cherry photo in the natural scene category) could be rated as most and least attractiveness depending on individual choice.

Thus, in Experiment 2, all the face and natural scene images were in equal luminance with each other as well as the background. Results showed that the amount of pupil constriction was linearly correlated with the attractiveness rating not only for faces [*F*(1,14) = 20.36, *p* < .001, Figure 2A and 2E] but for natural scenes as well [*F*(1,14) = 6.48, *p* < .03, Figure 2B and 2F]. When participants performed the roundness task, pupil constriction was still linearly correlated with the attractiveness of the faces [*F*(1,14) = 10.33, *p* < .01, Figure 2D and 2H], but not with the roundness judgment [*F*(1,14) = 3.43, *p* = .09, Figure 2C and 2G]. In summary, Experiment 2 replicated the main finding of pupil constriction to attractiveness faces and extended it to natural scenes. We can therefore conclude that pupil constriction certainly reflects attractiveness either explicitly or implicitly.

### Potential factors that may affect the effect of pupil constriction to attractiveness

Before we move on to the second focus of this study, namely whether pupil constriction also contributes to attractiveness judgment, we need to mention several additional, yet critical, issues. First, the biggest question would be why there is such a big inconsistency between our finding (pupil constriction to attractiveness) and the pupil dilation to attractiveness demonstrated in the literature. Second, it is still unclear whether the attractiveness judgment for geometric figures could indeed not induce corresponding pupil constriction. Finally, to what extent can the result be explained by gaze and/or eye accommodation factors? To address the first and second issues, we conducted three additional experiments (Experiments 3, 4 and 5) to examine the factors including stimulus presentation time, task demand, stimulus category, and overall pupil response pattern (constriction or dilation) caused by sequential luminance contrast change. In Experiment 3 when prolonging stimulus presentation time, pupil constriction to attractive faces was found not only during 0.5–1.5 s [*F*(1,16) = 18.98, *p* < .001] but also during 3–5 s [*F*(1,16) = 9.41, *p* < .01] after the stimulus onset (Figure 3). In Experiment 4 when using line-drawing faces with different task demand, the overall pattern of results changed (Figure 4). In the attractiveness rating condition, there was a tendency of early pupil constriction to attractiveness whereas it did not reach statistical significance [*F*(1,11) = 1.69, *p* = .2]. Most importantly, in contrast with the lasting effect of pupil constriction to attractiveness found in Experiment 3, the most attractive faces (rated as 8 or 9) induced the strongest pupil dilation response [*F*(6,66) = 3.10, *p* < .01] during the later time course (3–5 s). In the passive-viewing condition, although none of the effects were significant, there was a tendency of pupil dilation to the most attractive faces during the early time course [*F*(6,66) = 2.21, *p* = .05]. In Experiment 5, not only faces but also geometric figures were used to examine whether the general pupil response pattern (constriction or dilation) affects the effect. Results are shown in Figure 5. When the pupil in general dilated to faces, mean pupil size negatively correlated with attractiveness rating (i.e., the smaller the pupil, the more attractive the face was) although the effect was not significant [*F*(1,9) = 1.29, *p* = .3]. When the pupil in general constricted to geometric figures, there was still no tendency of correlation between pupil constriction and attractiveness [*F*(1,9) = 0.82, *p* = .4], consistent with the result in Experiment 1.

In sum, the overall results in Experiments 3-5 indicated that none of the factors that we examined alone can explain the discrepancy between our finding and the literature, but, in general, the effect of pupil constriction to attractive faces (still not to geometric figures) was more effectively observed during the early time course (within 2 s after the stimulus presentation, see Experiments 3 and 4), when the pupil generally constricted in response to the relative sequential luminance increase (compare Experiments 1 and 5) and when a task demand was required in contrast with passive viewing (see Experiment 4). On the other hand, pupil dilation to attractiveness was occasionally observed only for faces that were rated as most attractive (an 8 or 9 rating score), either during a later time period (3 to 5 s after stimulus onset, see Experiment 3) or passive viewing (Experiment 4). This is in a way consistent with the literature, where most of the evidence for pupil dilation to attractiveness came from pupillary responses accumulated for 10 s while participants just passively viewed the stimulus (e.g., Hess, 1965; Nunnally et al., 1967; Stass and Willis, 1967; Koff and Hawkes, 1968; Barlow, 1969; Atwood and Howell, 1971; Rieger and Savin-Williams, 2012).

In the following sections, we addressed the issue whether gaze and/or eye accommodation factors might explain the finding of pupil constriction to attractiveness, by conducting additional gaze-related analyses for Experiments 1-5. The results and figures are shown in Table 3 and Figures 12-14. First of all, we checked the gaze location data to examine where exactly the participant scrutinized on the target image. The heat maps of gaze location distribution during target presentation, superimposed on the sampled target image, are shown in Figure 12. As illustrated, the participant mostly looked at the center of the image in all Experiments 1-5. There was 80% of the gaze which was located within 3.7 degree of visual angle in eccentricity.

**Figure 12.**
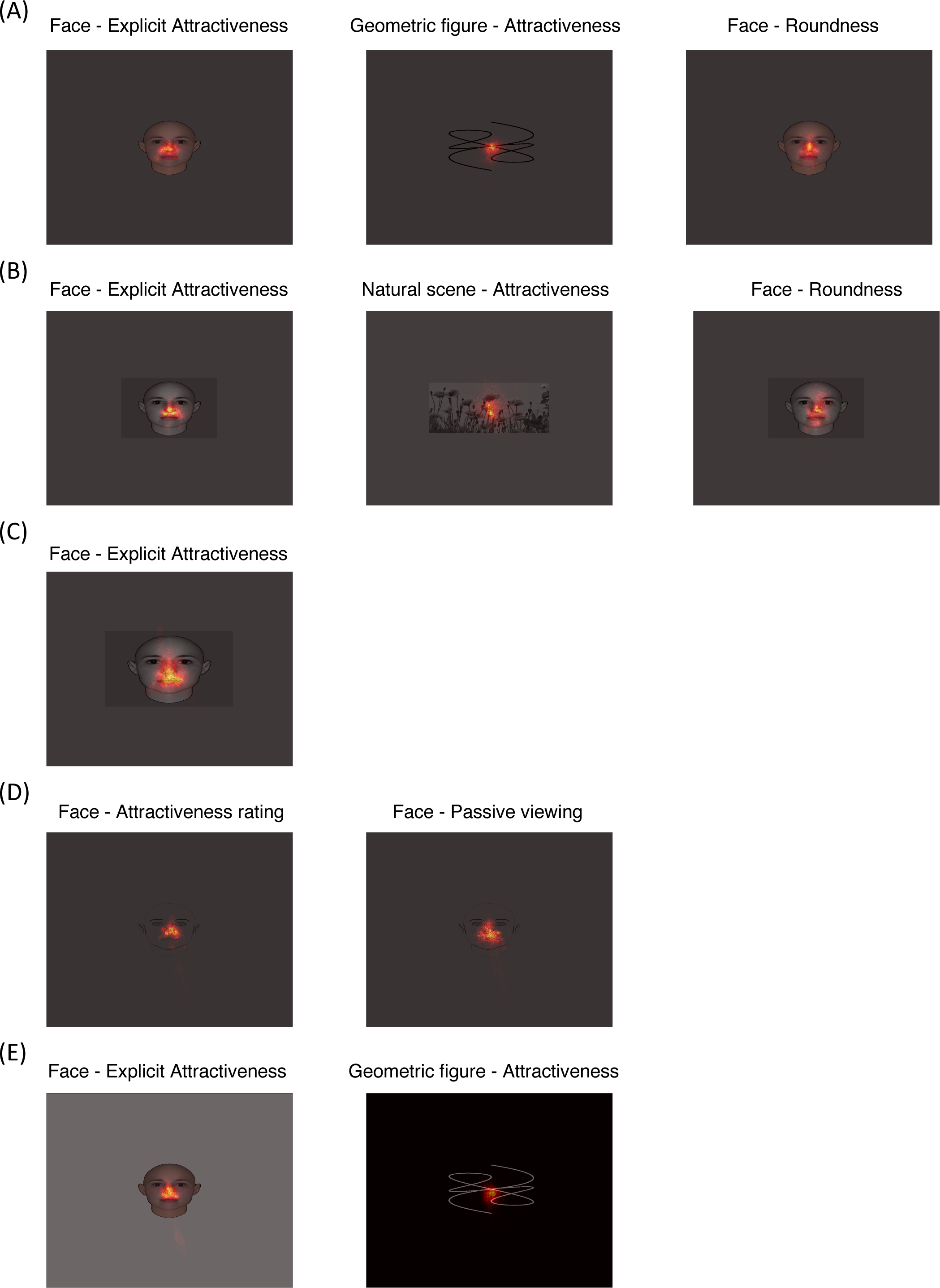
Heat map of gaze distribution during target presentation in Experiments 1 (A), 2 (B), 3 (C), 4 (D), and 5 (E), superimposed on the sampled target image. The target images were scaled to the visual display viewed in the real experiments. The luminance of the background represents approximate luminance in the real experiments: gray in (A)-(D), white in the face condition and black in the geometric figure condition in (E). The gaze data were accumulated from all trials and all participants.

**Figure 13.**
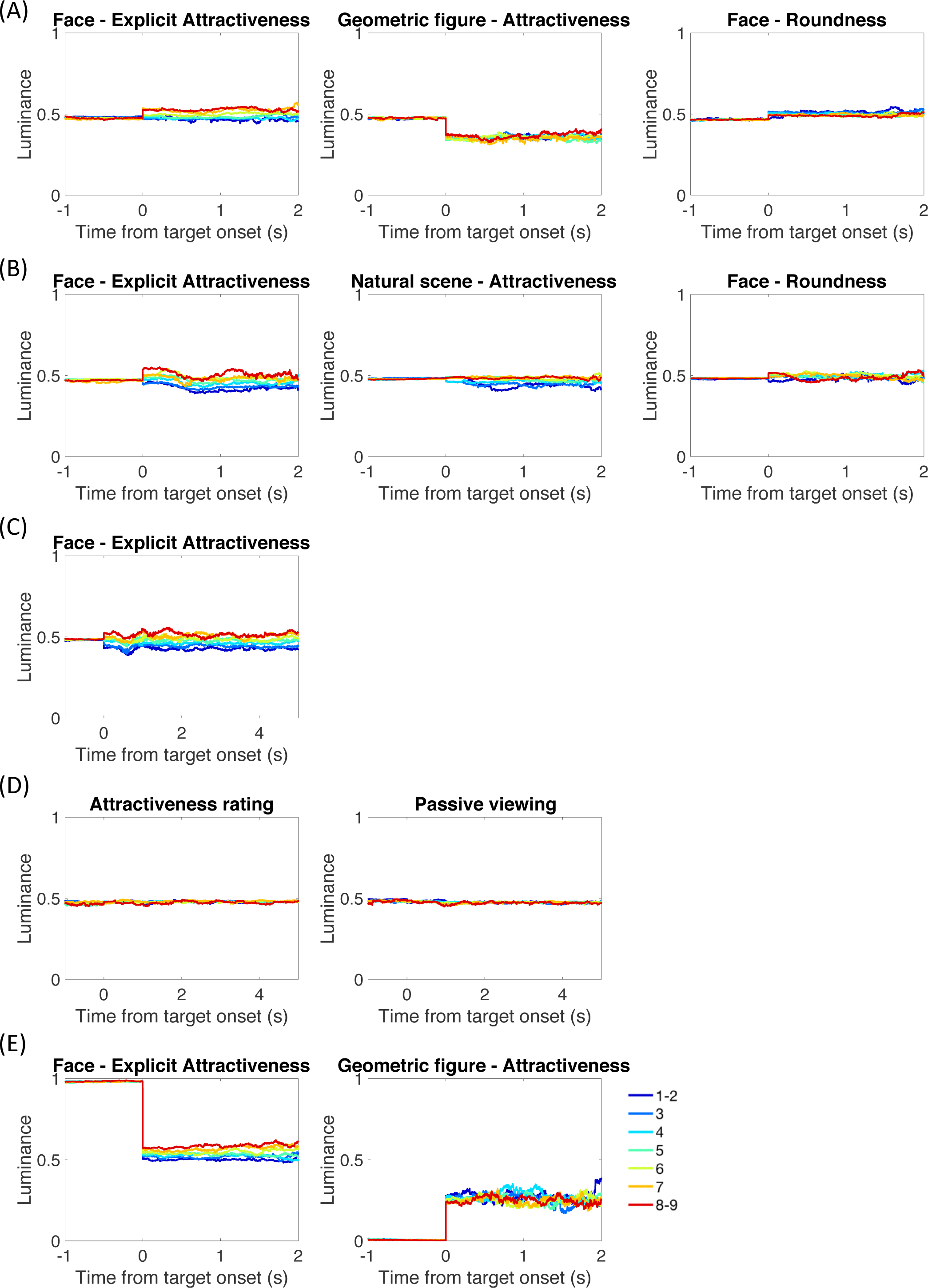
Mean local luminance contrast change as a function of time reference to the stimulus onset in Experiments 1 (A), 2 (B), 3 (C), 4 (D), and 5 (E). Local luminance contrast was calculated by averaging the luminance of the image region being gazed at, i.e., within1 degree of visual angle of the gazing point. Curves are parameterized with rating score depending on individual participants’ choices.

**Figure 14.**
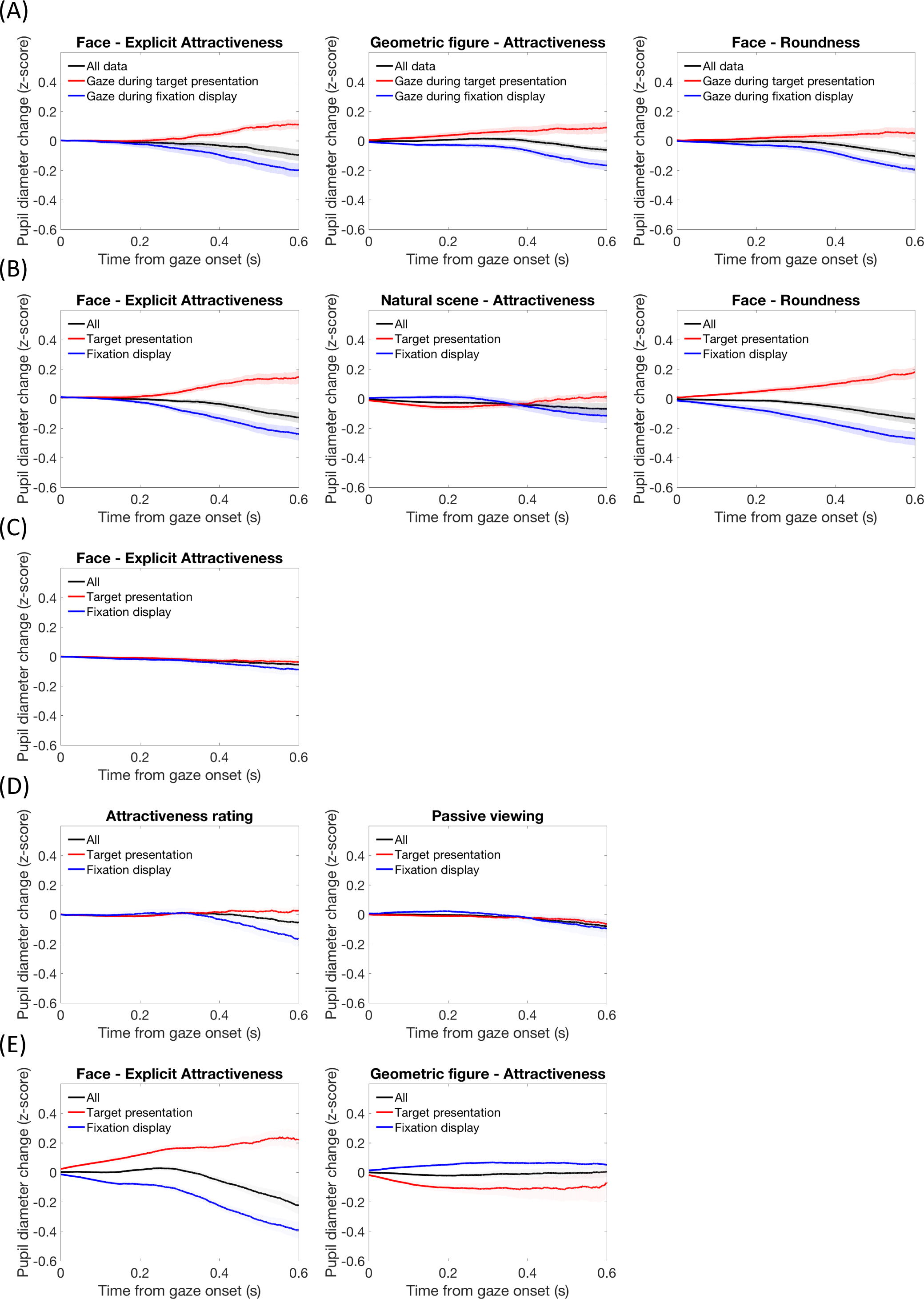
Gaze-contingent pupillary response in Experiments 1 (A), 2 (B), 3 (C), 4 (D), and 5 (E). Mean pupil diameter change as a function of time reference to the gaze onset, parameterized by the gaze being detected during the period of target presentation or fixation display.

While the gaze location was mainly distributed at the center of the target image and the average luminance of the images were equated in Experiments 2, 3, and 4, there might be still subtle luminance differences of the image regions being gazed at, and this might explain why pupil constricted stronger to attractive images.

To clarify this concern, we calculated the luminance of the image regions being gazed at, parameterized by rating score (see Figure 13). Result showed that participants tended to prefer the face image (but not the natural scene images or line drawing faces) with higher local luminance contrast in the center. To further examine whether the local luminance (of the image regions being gazed at) might explain the finding of pupil constriction reflecting attractiveness rating, we conducted a partial correlation analysis in which the correlation between mean pupil size and attractiveness rating was estimated when the effect of mean gazed local luminance was removed. Results are shown in Table 3. As expected, the correlation between pupil size and attractiveness rating was indeed reduced in certain conditions. Importantly, the correlation was still significant in most critical conditions: the face-implicit attractiveness condition in Experiment 1 (*p* < .05), the face-explicit attractiveness and natural scene-attractiveness conditions in Experiment 2 (*p*s < .03), and face-explicit attractiveness condition in Experiment 3 (*p* < .01). The overall results are consistent with our conclusion in Experiment 2 that early transient pupil constriction reflects attractiveness when the image luminance is controlled.

Accommodation to focus on near objects also induces pupil constriction, i.e., pupil near response (Mays and Gamlin, 1995; McDougal and Gamlin, 2008). A parsimonious hypothesis why pupil constriction reflects attractiveness would be due to a co-occurrence of eye accommodation and pupil constriction, assuming that gazing and/or focusing alone explains the effect. To test this hypothesis, we examined how the pupil responded during accommodation when viewing attractive visual objects. If pupil constricted near focus time and this pupil constriction following the focus to the visual object explained pupil constriction to attractiveness, we expected to observe this focus-evoked pupil constriction more vigorously when seeing attractive visual objects.

We conducted the gaze-contingent pupillary response analysis in which we aligned the pupil size data with the gaze onset. Results are shown in Figure 14. In accordance with the pupil near response, pupils on average constricted following the gaze (i.e., during accommodation) when viewing fixation display (the blue lines in Figure 14, except for the geometric figure condition in Experiment 5 since the luminance of the background was black whereas it was white or gray in all the other conditions). Most importantly, however, pupils in general dilated, instead of constricted, following the gaze when viewing target images (the red lines in Figure 14). This result surprised us in a sense that pupils were expected to constrict near focus time, in particular when the face image was brighter than the background, e.g., in Experiment 1. We suspect that the pupil dilation could be triggered by task demand, motor commend, and/or emotional arousal to the faces as the dilation was less observed when the rating response was postponed (Experiments 3 and 4), when the participant viewed the images without any task (the passive viewing condition in Experiment 4), or when the to-be-judged stimuli were natural scenes (Experiment 2) or geometric figures (Experiment 1). In any case, the accommodation to focus on the visual object did not necessarily lead to pupil constriction, at least under the circumstance in our experimental setup.

Together with the findings that the overall amount of pupil constriction reflected attractiveness judgment regardless of whether the rating response was asked immediately (Experiments 1 and 2) or postpone (Experiment 3), and whether the visual objects were faces or natural scenes, the overall results indicate a clear dissociation between pupillary responses to accommodation and to attractiveness. In other words, the gaze-contingent pupillary response result indicates that the eye accommodation alone cannot explain the finding of pupil constriction to attractiveness.

### Positive loop between pupil constriction and attractiveness

Now, the second main objective of the current study was to examine whether pupil constriction contributes to facial attractiveness judgment. In Experiment 6, the amount of pupil constriction was manipulated by relative background luminance changes (from the pre-stimulus screen) to examine whether the attractiveness judgment for faces changed accordingly. As shown in Figure 6B, displaying the face after the black fixation display caused stronger pupil constriction than displaying it after the gray fixation display did [*F*(1,10) = 278.43, *p* < .001]. The target background also affected the pupillary response in that the pupil constricted less for the black target background than it did for gray one [*F*(1,10) = 153.51, *p* < .001], whereas the influence of the pupil constriction was affected more strongly by the luminance of the fixation display than that of the target background [interaction: *F*(1,10) = 55.26, *p* < .001]. In a casual survey after the experiment, most participants reported that they were aware of the luminance change in the target background but mentioned little about the pre-stimulus display.

Critically, the attractiveness rating results are consistent with our hypothesis that when the pupil constricts more, the face is evaluated as more attractive (see Figure 6B and 6C for individual data). Specifically, parallel to the amounts of pupil constriction, faces were rated more attractive following the black fixation display [mean rating score = 4.63 vs. 4.40, *F*(1,10) = 7.22, *p* < .03]. Note that the face images were exactly the same in their identities as well as in their luminance in both the black and grey fixation display conditions (see Materials and Methods for details). The results can only be attributed to the pre-stimulus background luminance changes. In contrast, the target background by itself did not affect the rating [mean rating scores of 4.53 and 4.50 for the black and gray target background, respectively; *F*(1,10) = 0.18, *p* = .68]. The non-significant difference in the rating between the two types of target background indicated that the simultaneous contrast alone could not affect facial attractiveness judgments. Although the target background also induced significant changes in pupil size, the effect may interact with target background luminance itself to obscure its influence on attractiveness judgment. This is consistent with the casual survey in that some participants claimed that the target background might have affected their attractiveness judgment, but how it might have done so was not consistent among their reports. Alternatively, the non-observed effect of target background to attractiveness judgment may be due to the weaker modulation of the pupil constriction compared to the effect induced by the pre-stimulus display. Either way, the results are consistent overall with the interpretation that pupil constriction due to the sequential luminance contrast shift (from the pre-stimulus background to the stimulus) leads to higher ratings of attractiveness, and that in most cases, people are not aware of the causal relationship there.

We would like to emphasize that in our hypothesis, it is the amount of pupil constriction (i.e., pupil constriction amplitude), but not the pre-stimulus luminance, that contributes to attraction. In order to directly investigate the causal relationship among pupil constriction amplitude, pre-stimulus luminance (associated with pre-stimulus pupil baseline), and attractiveness rating, we conducted partial correlation analyses to examine whether it was the pupil constriction amplitude or pre-stimulus pupil baseline that predicted attractiveness rating when the effect of the other pupil-related factor was removed. Results showed that, consistent with our hypothesis, there was a significant correlation between pupil constriction amplitude and attractiveness rating when the effect of pre-stimulus pupil baseline was removed [mean *r* = -0.07, *t*(10) = -5.61, *p* < .0001]. By contrast, the correlation between pre-stimulus pupil baseline and attractiveness rating was not significant when the effect of pupil constriction amplitude was removed [mean *r* = -0.02, *t*(10) = -0.77, *p* = .46]. The overall results indicated a direct link between pupil constriction amplitude, but not pre-stimulus pupil baseline (or pre-stimulus luminance), and attractiveness rating.

However, one may still argue that either adaptation to the pre-stimulus luminance (i.e., the fixation display’s luminance) or sequential contrast may lead to brightness differences in faces, which may affect the attractiveness judgment. In Experiment 7, we examined whether sequential luminance contrast alone, when not inducing a strong difference in pupil response, causes differences in attractiveness judgments. Here the sequential luminance contrast is defined as Weber contrast, i.e., (*I* – *I*b) / *I*b with *I* and *I*b representing the luminance of the target images and the background, where only the local region surrounded the target image is taking into account. As shown in the experimental procedure (Figure 7A), the (local) sequential luminance contrast was changed between the conditions when the face followed the black and white hemifield. By contrast, the pupil response was expected to be similar between the two conditions assuming that pupil responded to overall global luminance change in general.

Results are shown in Figure 7B and 7C. As predicted, the pupil constricted similarly regardless of whether the face image followed the black or white hemifield in the pre-stimulus fixation display, although the pupil constriction difference was (marginally) significant in opposite patterns depending on the eye movement condition [the overall gaze location was confirmed to be on the face image in the peripheral in Experiment 7a and to be at the central fixation point in Experiment 7b, see Figure 15; the pupil constricted more strongly when the face followed the white hemifield than it did when the face followed the black one in Experiment 7a, *t*(15) = 2.90, *p* = .01 and vice versa in Experiment 7b, *t*(16) = 2.12, *p* = .05]. Accordingly, facial attractiveness judgments showed similar scores between the two pre-stimulus local luminance conditions [*t*(15) = 0.59, *p* = .56 in Experiment 7a and *t*(16) = 1.63, *p* = .12 in Experiment 7b, see Figure 7D and 7E for individual data], while the slight rating difference tendency was in accordance with the amount of pupil constriction rather than the local sequential luminance contrast. The overall results suggest the rating differences found in Experiment 6 cannot be explained solely by the local sequential luminance contrast. Instead, it is more in accordance with the causal contribution of pupil constriction to attractiveness judgment.

**Figure 15.**
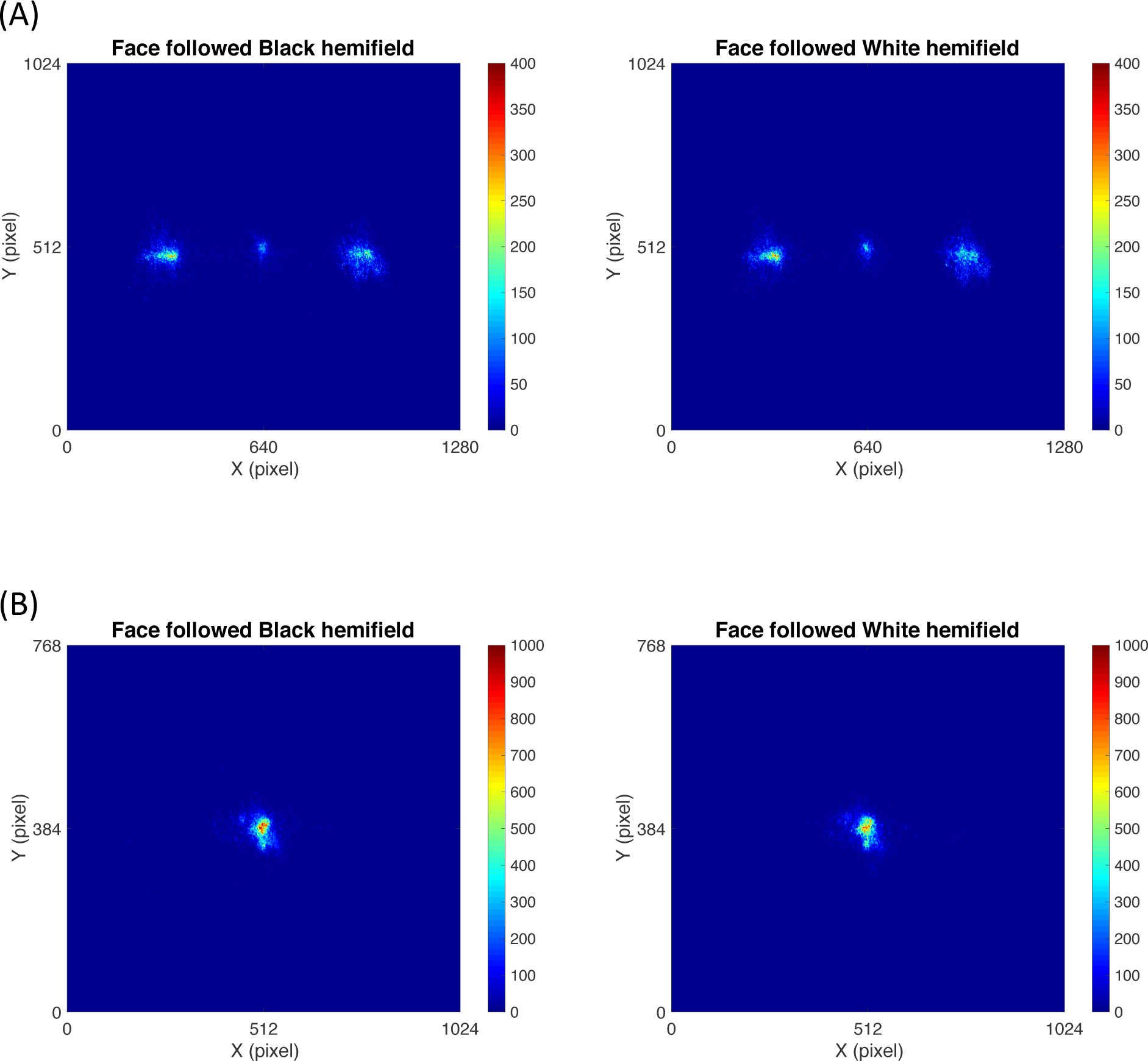
Heat map of gaze distribution during 0-2 s reference to the target onset in Experiment 7 when (A) eye movement was allowed or (B) eye movement was not allowed (participants fixated the central fixation cross throughout the trial).

## Discussion

We found an early and transient pupil constriction response in proportion to attractiveness judgment. The constriction response was found be specific to aesthetic object categories such as faces or natural scenes, as opposed to relative emotionally neutral objects such as line-drawing geometric figures. When participants were asked to judge the roundness of faces, pupil constriction still correlated with their attractiveness but not the roundness rating score, indicating the automaticity of the pupil constriction to attractiveness. The result of pupil constriction to attractive faces was replicated in three experiments by using various face stimuli: natural color images (Experiment 1) and equal luminance images (Experiments 2 and 3). Potential confounding factors such as gaze location and eye accommodation were ruled out. Moreover, when we manipulated pupillary responses implicitly by manipulating relative background luminance changes (from the pre-stimulus screen), the facial attractiveness ratings were in accordance with the amount of pupil constriction (Experiment 6), and the result could not be explained solely by simultaneous or sequential luminance contrast (Experiments 7a and 7b). In summary, we found a tight link between pupil constriction and facial attractiveness, at least under certain conditions. The finding could have profound implications with respect to the well-known theories in the James-Lange tradition concerning mind-body interaction. It could also provide new clues to the neuro-physiological mechanisms underlying attractiveness decisions and to implementing various real-world applications such as BMIs (brain-machine interfaces) and marketing strategies.

### Pupil constriction vs. dilation to attractiveness

Our counterintuitive finding of pupil constriction, rather than dilation, to facial attractiveness reveals a heretofore unknown relationship between the pupillary response and affective decision-making. The discrepancy between our results and those in the literature can be understood by considering three potential factors. First, the pupil is highly sensitive to subtle luminance (contrast) differences, and early studies (especially in the 60s and 70s) might be just not technically capable of controlling it. Second, the temporal scale of pupil size measurement may have been different in those classical studies relative to ours. Indeed, we found two phases of pupil response over time: early constriction (approximately 0 to 2 s from the stimulus onset), and late dilation (after 3 s) to attractive faces (Experiment 4). In contrast to our approach that analyzed the dynamic changes in the pupillary response on a finer scale, previous studies just took the average pupil size over time, typically 10 s after the stimulus presentation (e.g., Hess, 1965; Nunnally et al., 1967; Stass and Willis, 1967; Koff and Hawkes, 1968; Barlow, 1969; Atwood and Howell, 1971; Rieger and Savin-Williams, 2012). They may have glossed over the two dynamic phases of the pupil response (constriction, then dilation) and thus failed to reveal the early, transient component of the pupil constriction response to attractiveness judgment. Third, the cognitive state (mental set) has an influence on pupil response (e.g., Binda et al., 2013; Mathôt et al., 2013; Naber et al., 2013; Binda et al., 2014; Mathôt et al., 2014). The non-controlled cognitive state (i.e., passive viewing) in those classical studies (e.g., Hess, 1965; Nunnally et al., 1967; Stass and Willis, 1967; Koff and Hawkes, 1968; Barlow, 1969; Atwood and Howell, 1971; Rieger and Savin-Williams, 2012) made it difficult to uncover the effects of cognitive processes for decision and/or attractiveness evaluation per se. The involvement of cognitive processes may be deeper than just serving to reveal a different aspect of the relationship between the pupil response and attractiveness.

Putting the above factors aside, reviews of recent studies that demonstrate the correlation between pupil dilation and attractiveness judgment have revealed that the correlation really depends on the observer’s and the observed face’s gender (Simms, 1967; Rieger and Savin-Williams, 2012) and emotion (Harrison et al., 2006; Harrison et al., 2009), suggesting a more complicated mechanism than a straightforward linkage between pupil dilation and attractiveness. One must conclude the traditional belief that pupil dilation reflects attractedness simply does not account for what is really going on between the brain and the eyes. It is important for future studies to closely examine the dynamic changes in pupillary response over time to isolate the effects of cognitive processes and emotional arousal.

### Possible neural mechanism of pupil response to attractiveness

Pupil size is controlled by two sets of antagonistic muscles, the iris sphincter muscle and iris dilator muscle, innervated by parasympathetic and sympathetic nerves, respectively. It is thus naturally presumed that one possible underlying neural mechanism of the correlation between facial attractiveness and pupil constriction is based on the activation of the parasympathetic nervous system. Usui and Hirata (1995) proposed a nonlinear dynamical model for the human pupillary muscle plant. The model states that the human pupil response to a flash visual stimulus can be explained by a combination of an early, transient parasympathetic activation (within 2 s) and a slow, sustained deactivation of the sympathetic activation, and this was confirmed by pharmaceutical manipulation (Yamaji et al., 2000). This is consistent with the hypothesis that early, transient pupil constriction to attractiveness is driven by the parasympathetic nervous system. The cause of pupil dilation to attractiveness, in contrast, is more complicated. It could be due to emotional arousal activating the sympathetic nervous system (Bradley et al., 2008) during the longer time course and/or under a passive viewing situation, in particular when the pupil response is less affected by a flash visual stimulus. Alternatively, it could to due a deactivation and/or rebound of the parasympathetic nervous system following the early 2-s transient activation. In addition, other factors, such as stimulus properties and/or task demands, may activate the autonomic nervous system interactively. For instance, it is possible that the face and natural scene images used in the current study are more likely to induce a joyful, relaxing, and/or soothing experience that actives the parasympathetic nervous system, in contrast to inducing excitement, which may active the sympathetic nervous system dominantly. In the same vein, the failure of observing any effect with geometric figures may be due to the lack of attractiveness of the images we used in general, or the high similarity of the images which made the attractiveness judgment less differentiable, compared with faces or natural scenes. It is conceivable that attractiveness has multiple meanings, and the judgment may change depending on the context (e.g., when choosing a life partner vs. a queen in a beauty contest). The relationship between pupil response and attractiveness is not as simple as conventionally believed. To better understand the neural mechanism of attractiveness formation, further studies should investigate how different factors such as a stimulus’s emotional valance and strength affect its attractiveness, together with other physiological measurements under different time courses. This approach may have potential impact on decoding complicated emotions instigated by the interaction between the sympathetic and parasympathetic nervous systems from eye metrics. In any case, our finding of pupil constriction to attractiveness, after eliminating various artifacts/side factors, is sufficient to raise the warning, in the least.

That said, this parasympathetic nervous system hypothesis alone does not directly account for the causal contribution of pupil constriction to facial attractiveness. Instead, one may need to assume some sort of positive loop between liking and seeing to understand all the results that we report here. According to the positive loop account, the longer we see, the more we like, and vice versa, which is supported further by Shimojo et al.’s earlier findings of the gaze cascade (Shimojo et al., 2003). The pupil constricts to increase visual acuity/clarity to obtain a sharper facial image, to make it more attractive, or to facilitate prolonged inspection time. Moreover, prolonged inspection may further activate the parasympathetic nervous system, by accompanying the feeling of calming and soothing. These may together participate in the decision-making formation of liking. While highly speculative, this scenario is not only feasible physiologically but nicely incorporates the parasympathetic account as a part of an entire dynamic loop as well, and is thus consistent with both the correlation results (Experiments 1, 2 and 3) and the causal results (Experiment 6). Further studies using different approaches to manipulate the pupils are required to examine our hypothesis and/or investigate the underlying neural mechanisms. For instances, one may use eye-drop-administered cholinergic drug to manipulate the pupil size at different timing reference to the stimulus presentation to investigate how quickly or slowly the autonomic nervous system interacts with the eye-controlled system. Alternatively, one may use artificial pupils (e.g., a piece of paper with a tiny hole) to adjust the amount of image information into retina to investigate the relative contribution of the retinal input and physiological/muscular control of the pupil size to attractiveness.

### Mind-body interaction and implications

The finding that the pupil manipulation contributes to facial attractiveness judgments should be added to the long list of evidence for the James-Lange tradition of body-mind causality, regardless of whether the above parasympathetic account and positive-loop interpretations are valid. In addition to the classical association between physiological arousal and experienced emotion such as euphoria and anger (Schachter, 1964), our findings reveal an until now unknown physiological cause, i.e., pupil constriction, to mind (facial attractiveness judgment). While the physiological status is altered for unknown reasons, a reason has to be given at the conscious level. This is not that surprising as shown in the suspension bridge effect (Dutton and Aron, 1974), where people tend to misattribute unknown physiological arousal, i.e., the anxiety induced by walking on a suspension bridge, to romantic attraction. In our case, the physiological change, i.e., the pupil constriction, is (mis)attributed to evaluative attitudes towards facial attractiveness. The possible prolonged looking behavior due to the pupil constriction response is (mis)attributed to the preference for the seen image.

In the same vein, our finding can be also interpreted as a new example of “cognitive dissonance” and its solution, i.e., pupil constriction, at the implicit level.

Cognitive dissonance refers to a mental state where a person holds more than two contradictory beliefs, ideas, or attitudes at the same time and experiences uncomfortable stress because of that. In relation to affective decision-making, it has been shown that choice per se creates preference for the chosen object (Brehm, 1956) to reduce cognitive dissonance (i.e., people would not choose an object which they do not like). The same logic also applies to how inspection per se affects preference during which the brain and eyes, including gaze and pupil response, are involved. People prefer the object that they look at longer (Zajonc, 1968; Shimojo et al., 2003). In contrast to gaze, the contribution of the pupil response to decision-making is implicit in two senses. First, it is an automatic response that is nearly impossible to voluntarily control. Second, the process of facial attractiveness formation via pupil constriction is hardly identifiable by attentive causal introspection.

After decades of neglect, pupillometry has been recently been revived by studies showing that pupil response reflects various cognitive processes, including attention (Aston-Jones and Cohen, 2005; Einhäuser et al., 2008; Eldar et al., 2013), memory (Goldinger and Papesh, 2012; Naber et al., 2013; Zokaei et al., 2019), decision making (Einhäuser et al., 2010; de Gee et al., 2014), and linguistic (Schmidtke, 2018) and auditory processing (Liao et al., 2016a; Liao et al., 2016b; Zhao et al., 2019). However, re-examining its relationship with attractiveness judgment has attracted little interest, because of the belief in the correlation between pupil dilation and attractiveness. The current study uncovers a heretofore unknown tight link between pupil constriction and attractiveness. Additionally, it also indicates that pupil response likely participates in the mechanism underlying attractiveness judgment formation. Our finding goes beyond the scope of reading the mind from the eyes, to further imply that the neural mechanism that controls pupil responses also massively interacts with higher-level cognitive processes such as preference formation.

